# RPRM (Reprimo) triggers SCF^FBXW11^-mediated DNA-PKcs degradation to block non-homologous end joining and radiosensitize tumors

**DOI:** 10.64898/2025.12.14.694167

**Authors:** Zhaojun Li, Yarui Zhang, Binghua Tang, Shuning Xu, Zhiyan Zhu, Jingdong Wang, Guomin Ou, Jinsheng Hong, Minglian Qiu, Hongying Yang

**Author notes:** Correspondence: M Qiu; H. Yang. These authors contributed equally to this work (Zhaojun Li, Yarui Zhang, Binghua Tang).

## Abstract

Targeting DNA damage repair pathways represents a promising strategy in cancer radiotherapy, however, the limited insight into the regulation of the non-homologous end joining (NHEJ) repair pathway in cancer cells severely restricts the development of precise radiosensitization approaches. Here, we identify Reprimo (RPRM), a p53-inducible tumor suppressor, is a potent inhibitor of NHEJ in which RPRM promotes proteasomal degradation of DNA-dependent protein kinase catalytic subunit (DNA-PKcs) phosphorylated at the T2609 site. Upon irradiation, RPRM quickly translocates into the nucleus to enhance the transcription of *FBXW11*, a F-box protein gene that encodes the substrate recognition component of SCF (SKP1-CUL1-F-box protein) E3 ubiquitin-protein ligase complex. In the nucleus, RPRM is also able to scaffold the assembly of the SCF^FBXW11^ E3 ligase complex by binding to both CUL1 and FBXW11. This complex can specifically ubiquitylate phosphorylated DNA-PKcs (T2609), leading to reduced levels of both phosphorylated DNA-PKcs (T2609) and total DNA-PKcs, thereby suppressing NHEJ-driven repair and enhancing tumor radiosensitivity. In consistence, inhibiting the SCF^FBXW11^ E3 ligase complex eliminates RPRM-mediated radiosensitization by protecting DNA-PKcs. Clinically, high *RPRM* expression in non-small cell lung cancer (NSCLC) predicts better patient prognosis. Moreover, *RPRM* expression correlates positively with *FBXW11* and *PRKDC* (DNA-PKcs) in the tumor tissues of the NSCLC patient cohort. Significantly, high expression of RPRM protein is linked to enhanced radiosensitivity of patient-derived organoids (PDOs) from these NSCLC patients. These results establish RPRM as a key negative regulator in NHEJ repair pathway and the RPRM-SCF^FBXW11^-DNA-PKcs axis as a potential target to enhance cancer radiosensitivity.

## Introduction

Despite significant advances in cancer radiotherapy, treatment outcomes continue to be undermined by radioresistance of cancer cells. This phenomenon is largely mediated by the efficient activation of their DNA damage response (DDR) pathways, which are triggered by the induction of DNA double-strand breaks (DSBs)—the most lethal lesions caused by ionizing radiation (IR) [1,2]. To maintain genome integrity and cell viability, cancer cells undergo transient cell cycle arrest and repair DSBs primarily through two classical pathways: homologous recombination (HR) and non-homologous end joining (NHEJ). While HR requires a homologous template for precise repair and is thus restricted to the S/G2 phases of the cell cycle, NHEJ operates throughout the cell cycle and serves as the major DSB repair mechanism [3–5]. Therefore, to overcome radioresistance, a promising strategy is to target these DNA damage repair pathways especially the NHEJ, thereby sensitizing cancer cells to IR.

DNA-PKcs, a phosphatidylinositol 3-kinase-related kinase (PIKK), is well known as a classic non-homologous end joining factor. Following DSB induction and limited end-processing by the MRE11-RAD50-NBS1 (MRN) complex, DNA-PKcs is recruited to the DSB sites, where it forms the functional DNA-PK complex with the Ku70/80 heterodimer and other key proteins like Artemis and XRCC4-like factor (XLF). The complex then phosphorylates itself and other targets, controls DNA-end sensing, protection, processing and ligation in NHEJ [6–9]. Its core role in NHEJ provides a robust rationale for targeting DNA-PKcs in cancer radiotherapy and genotoxic chemotherapy, leading to intensive development of DNA-PKcs inhibitors, although the application of DNA-PKcs inhibitors has lagged behind that of ATM inhibitors mainly due to their suboptimal pharmacokinetic properties and significant off-target effects [10]. Among these inhibitors, several, such as AZD7648 and NU5455, have demonstrated significant potential in enhancing the efficacy of radiotherapy and chemotherapy in mouse models [11,12], some, including M3814 and AZD7648, have advanced to clinical trials [13–16]. However, only limited anti-cancer effects have been observed so far in these studies [14,16]. This underscores the need for further investigation into the regulatory mechanisms of DNA-PKcs. Although both the critical functions of DNA-PKcs in the DDR signaling network and its underlying mechanisms have been well established [6,7], knowledge of its upstream regulators remains incomplete. Little is known about the regulation of DNA-PKcs turnover, beyond the confirmed role of E3 ubiquitin ligases membrane-associated RING finger protein 5 (MARCH5), Cullin 4A (CUL4A) and RING finger protein 144A (RNF144A) in its ubiquitylation and degradation [17–19]. Additionally, GNB1L, a poorly-characterized protein, may regulate the levels of both ataxia-telangiectasia mutated (ATM, another PIKK member) and DNA-PKcs through unknown mechanisms [20]. We have recently identified Reprimo (RPRM) protein as a negative regulator of ATM [21]. *RPRM* gene, a target of p53, is a tumor suppressor that is involved in a variety of cancers including gastric, breast, esophageal, pancreatic and pituitary cancers [22–27]. Limited studies have revealed its regulatory effects on G2/M cell cycle arrest, cell proliferation, cancer cell migration and invasion [22,28–30]. RPRM can be induced by DNA damage, in turn promoting DNA damage-induced apoptosis, suggesting that it plays a role in DDR [21,22,31]. Our previous studies have demonstrated that RPRM is indeed a critical mediator of DNA damage repair and cellular radiosensitivity, thus, knockout of *Rprm* gene confers significant radioresistance in mice [21,32–34]. Furthermore, we have uncovered a mechanism underlying its regulation on DNA repair. Upon induction, RPRM protein translocates into the nucleus and binds to ATM, promoting its nuclear export and proteasomal degradation. This subsequently leads to impaired DNA repair, enhanced apoptotic cell death and increased sensitivity to DNA damage. More specifically, both HR and NHEJ pathways are inhibited by RPRM, suggesting that apart from ATM, RPRM may also regulate DNA-PKcs [21]. Coincidentally, we also found that *Rprm* knockout promotes DNA-PKcs activation in hematopoietic stem cells of mice [32]. However, whether RPRM is an important regulator of DNA-PKcs and the mechanism underlying this regulation remain to be elucidated.

In this study, we demonstrated that RPRM had a greater negative impact on the NHEJ pathway than on the HR pathway, and revealed its suppressive effect on DNA-PKcs. RPRM promotes ATM nuclear-cytoplasmic translocation and degradation, whereas here we discovered that DNA-PKcs degradation regulated by RPRM occurred in the nucleus through a completely different mechanism. We also evaluated the association between *RPRM* expression and clinical outcomes in NSCLC.

## Results

### RPRM, a novel prognostic biomarker, inhibits the NHEJ repair pathway more potently than the HR pathway

The tumor suppressor *RPRM* is implicated in various cancers, such as breast, gastric, pancreatic, pituitary cancers, etc [23–31]. However, the association between *RPRM* and lung cancer remains unclear. By analyzing *RPRM* expression in NSCLC using The Cancer Genome Atlas (TCGA) database, we observed heterogeneous expression of *RPRM* among patients, with levels exhibiting a broad spectrum ranging from negligible to high, which is similar to the *RPRM* expression pattern in breast and pancreatic cancer. However, it was noted that the *RPRM* expression in NSCLC demonstrated a more dispersed pattern without a distinct peak of concentration, whereas its expression in breast and pancreatic cancer was predominantly at low to moderate levels, exhibiting a skewed distribution (Figure S1A). Additionally, we evaluated both mRNA and protein expression of *RPRM* in 20 clinical samples from NSCLC patients (Table S1), which further confirmed its heterogeneous expression pattern (Figure S1B and S1C). Most importantly, analysis of the TCGA NSCLC dataset showed that high *RPRM* expression correlates with better patient prognosis, revealing a consistent pattern across breast and pancreatic cancer (Figure 1A). These results indicate that *RPRM* is a promising prognostic biomarker in cancer.

**Figure 1.**
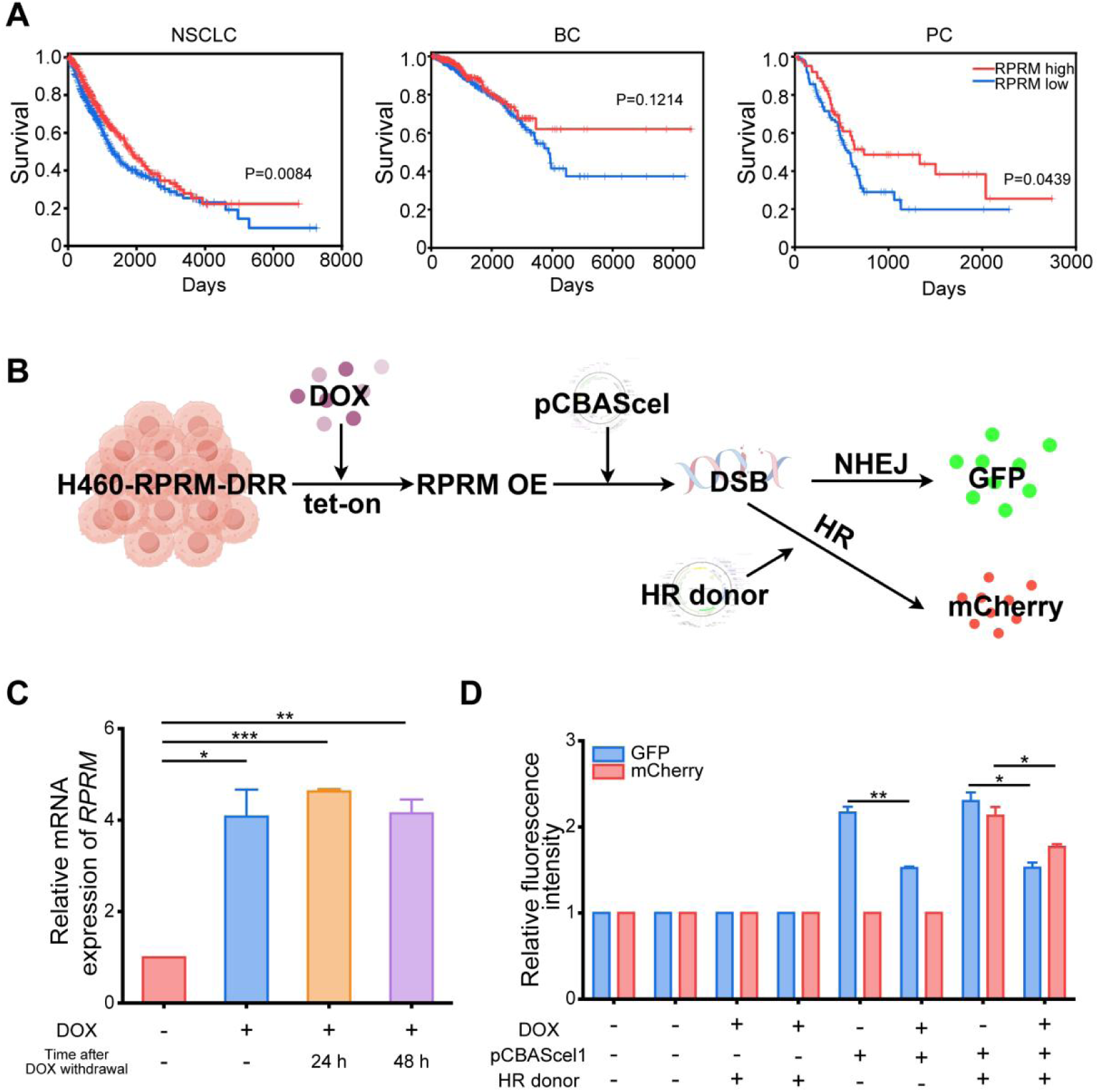
RPRM, a promising prognostic biomarker, inhibits NHEJ more potently than HR. A. Analysis of The TCGA dataset demonstrates that survival of NSCLC, breast cancer (BC) and pancreatic cancer (PC) patients was associated with *RPRM* expression. Kaplan-Meier survival curves and Log rank statistics are shown. B. Schematic diagram of the tet-on-based DRR system. C. The expression level of *RPRM* gene in H460-*RPRM*-DRR (tet-on) cells after DOX (0.3 μg/ml) treatment for 24 h. D. The relative fluorescence intensity of GFP and mCherry in H460-*RPRM*-DRR (tet-on) cells after NHEJ and HR repair of inverted ISce1 cuts with/without DOX treatment. \**P* < 0.05, \*\**P* < 0.01, \*\*\**P* < 0.001; significance was determined by two-way ANOVA followed by Tukey’s test.

Our previous analysis of the TCGA lung squamous cell carcinoma dataset also showed that high *RPRM* expression correlates with better prognosis in patients receiving radiotherapy. This therapeutic relevance of *RPRM* in cancer radiotherapy may stem from its critical role in DNA damage repair [21]. We have found that overexpression of *RPRM* in cancer cells inhibited the formation of 53BP1 and Rad51 foci shortly after radiation exposure, established markers for the NHEJ and HR repair pathways, respectively [35,36], indicating that RPRM inhibits both NHEJ and HR [21]. To determine which pathway was more strongly suppressed by *RPRM* overexpression, we established a tet-on-based DSB Repair Reporter system (DRR system) [37]. As shown in Figure 1B, following induction of *RPRM* overexpression by doxycycline (DOX), this DRR system allows the quantitative comparison of NHEJ versus HR in the same cells by measuring GFP and mCherry expression as reporters for NHEJ and HR repair of inverted ISce1 cuts, respectively. *RPRM* overexpression induced by DOX administration was confirmed using qRT-PCR (Figure 1C). Not surprisingly, compared to the untreated control, DOX-treated cells exhibited a marked reduction in the fluorescence intensity of both GFP and mCherry, indicating that *RPRM* overexpression inhibited both the NHEJ and HR repair pathways. More importantly, *RPRM* overexpression more potently inhibited NHEJ than HR (30% reduction in GFP intensity vs 17% reduction in mCherry intensity) (Figure 1D).

### RPRM reduces the expression level of DNA-PKcs through promoting its proteasomal degradation in the nucleus

We have previously revealed that RPRM promotes ATM degradation, resulting in impaired DNA repair and increased cellular radiosensitivity [21]. Since RPRM exhibited a more robust inhibition of NHEJ relative to HR (Figure 1), we hypothesized that the inhibition of the NHEJ repair pathway by RPRM was related not only to ATM [38] but also to DNA-PKcs, a key PIKK in this pathway [6,7]. Unexpectedly, *RPRM* overexpression alone promoted the transcription of *PRKDC* (Figure 2A); however, following 2 Gy X-irradiation, RPRM downregulated DNA-PKcs expression, with *RPRM*-overexpressing H460 cells (H460-*RPRM*) exhibiting significantly lower DNA-PKcs levels than the negative control cells (H460-NC) (Figure 2B). This result was also confirmed using immunofluorescence microscopy (Figure 2C). Moreover, this downregulation of DNA-PKcs by RPRM was consistently observed in both AGS and A549 cells (Figure S2), indicating that RPRM downregulates DNA-PKcs at the protein level.

**Figure 2.**
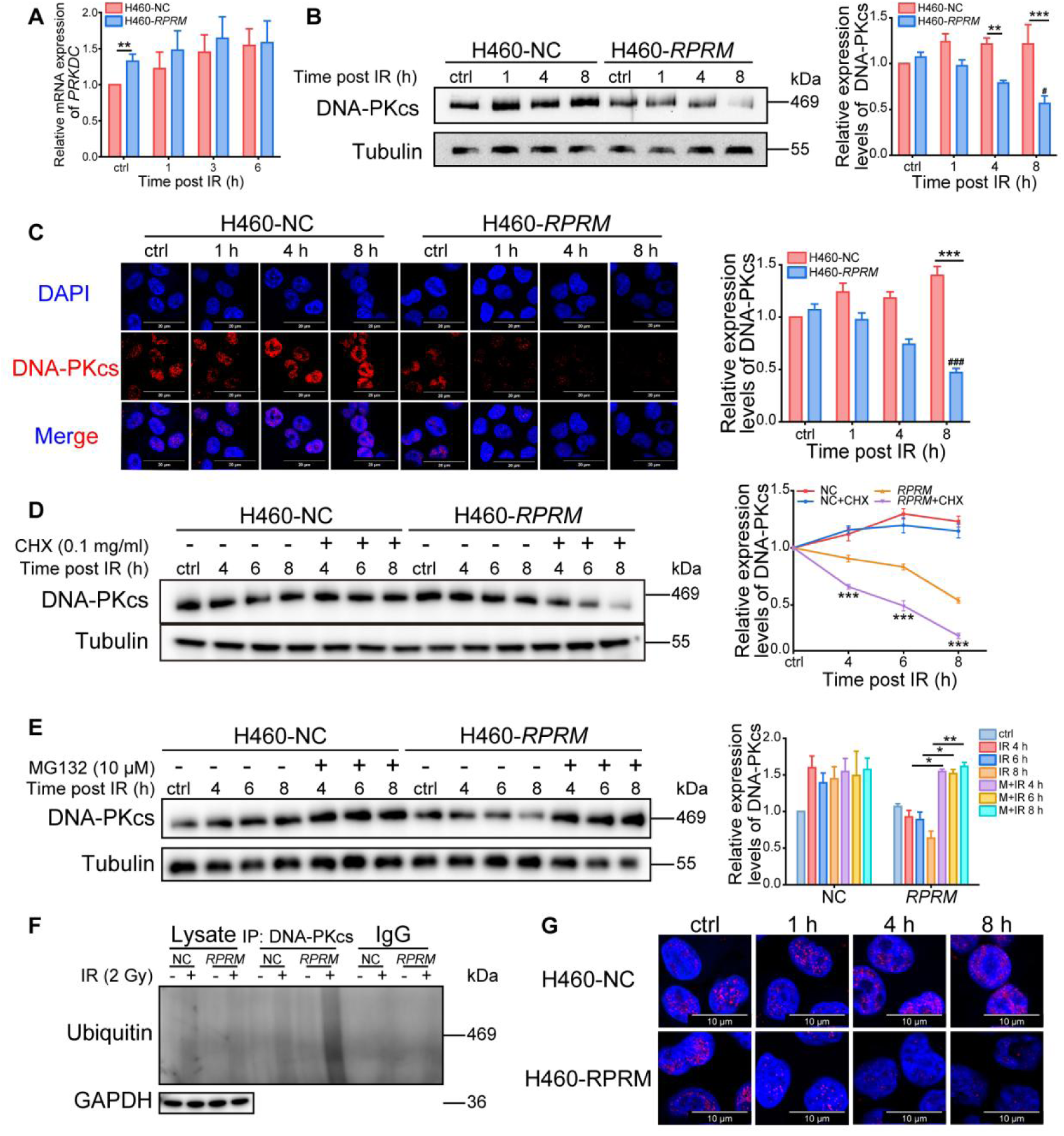
RPRM promotes DNA-PKcs degradation. A. The mRNA levels of *PRKDC* in H460-NC/*RPRM* cells after 2 Gy X-irradiation. B. The protein levels of DNA-PKcs in H460-NC/*RPRM* cells after 2 Gy X-irradiation. #, H460-*RPRM* IR-8 h vs H460-*RPRM* ctrl. C. Representative immunofluorescence images of DNA-PKcs in H460-NC/*RPRM* cells after 2 Gy X-irradiation and the quantification of the relative fluorescence intensity of DNA-PKcs. Scale bars, 20 μm. Two hundred cells were analyzed for each group. ###, H460-*RPRM* IR-8 h vs H460-*RPRM* ctrl. D. The protein levels of DNA-PKcs in H460-NC/*RPRM* cells after 2 Gy X-irradiation with/without CHX (0.1 mg/ml) pretreatment for 2 h. ***, *RPRM*+CHX IR vs *RPRM* IR. E. The protein levels of DNA-PKcs in H460-NC/*RPRM* cells after 2 Gy X-irradiation with/without MG132 (10 μM) pretreatment for 1 h. F. Change in DNA-PKcs ubiquitylation in H460-NC (NC) and H460-*RPRM* (*RPRM*) cells 4 h post 2 Gy X-irradiation detected by Co-IP. G. Enlarged immunofluorescence images of DNA-PKcs in H460-NC/RPRM cells after 2 Gy X-irradiation showing no obvious DNA-PKcs translocation. Scale bars, 10 μm. * and #*P* < 0.05, \*\**P* < 0.01, *** and ###*P* < 0.001; significance was determined by two-way ANOVA followed by Tukey’s test.

To investigate whether RPRM promotes DNA-PKcs degradation, cells were first treated with cycloheximide (CHX), a protein synthesis inhibitor, prior to irradiation. In contrast to H460-NC cells, DNA-PKcs levels in H460-*RPRM* cells was significantly reduced by this treatment following irradiation (Figure 2D). Conversely, inhibiting the proteasome with MG132 prior to irradiation significantly increased DNA-PKcs levels in H460-*RPRM* cells but not in H460-NC cells (Figure 2E). Additionally, following irradiation, DNA-PKcs ubiquitylation was significantly higher in H460-*RPRM* cells than in H460-NC cells (Figure 2F). These data indicate that RPRM promotes DNA-PKcs degradation by the ubiquitin-proteasome system (UPS) after X-irradiation.

Interestingly, RPRM facilitates the degradation of ATM by promoting its translocation from the nucleus to the cytoplasm, where ATM is degraded [21]. However, DNA-PKcs degradation in H460-*RPRM* cells following X-irradiation predominantly occurred in the nucleus without obvious nuclear-cytoplasmic translocation relative to H460-NC cells (Figure 2G). This suggests that RPRM regulates the degradation of DNA-PKcs and ATM through distinct mechanisms.

### RPRM promotes DNA-PKcs degradation through SCF^FBXW11^ complex

To explore the underlying mechanisms by which RPRM promotes DNA-PKcs degradation, we first determined whether RPRM interacted with DNA-PKcs. Surprisingly, RPRM, which binds to ATM [21], did not bind to DNA-PKcs either before or after irradiation (Figure 3A). Instead, RPRM interacted with both CUL1 and FBXW11, two components of SKP1-Cullin 1-F-box protein (SCF) complex (Figure 3B and S3). This finding confirms previous large-scale mass spectrometry study that reported an interaction between RPRM and these two proteins [39]. As a member of the cullin-RING E3 ligase (CRL) complex family, the largest E3 ligase family, SCF complex determines the substrate specificity for ubiquitylation and subsequent degradation [40,41]. Among the components of SCF complex, CUL1 is a scaffold protein that needs to be activated by conjugation with NEDD8, a small ubiquitin-like protein, for CRL function [41], and FBXW11 is an F-box protein that selectively recognizes and binds to specific substrate for ubiquitylation [40]. Therefore, we hypothesized that RPRM promoted DNA-PKcs degradation through SCF^FBXW11^ complex.

**Figure 3.**
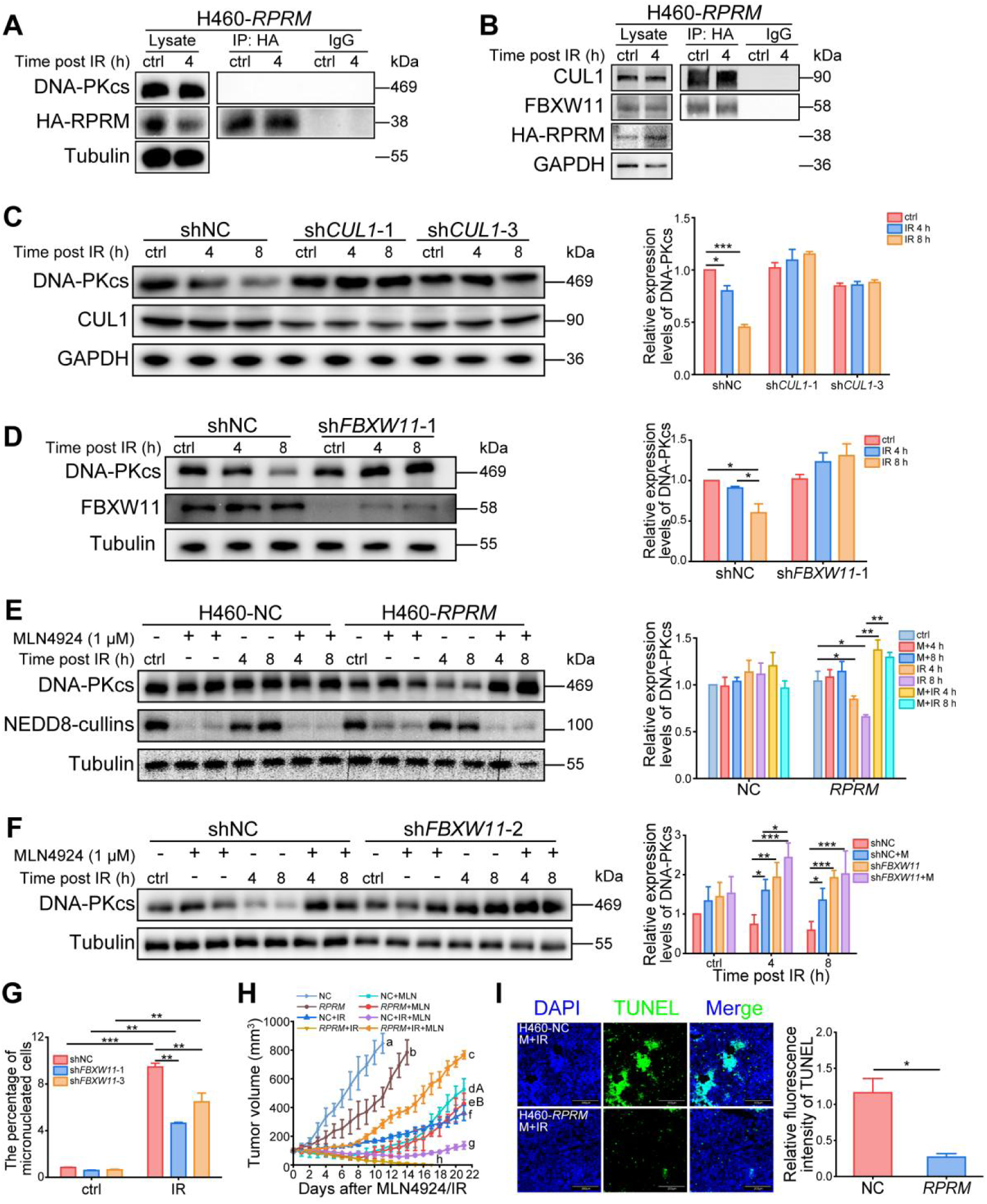
RPRM promotes DNA-PKcs degradation through SCF^FBXW11^ complex. A. No interaction between RPRM and DNA-PKcs was detected in non-irradiated and irradiated (2 Gy) H460-*RPRM* cells by Co-IP. B. Co-IP demonstrating that RPRM bound to both CUL1 and FBXW11 in non-irradiated and irradiated (2 Gy) H460-*RPRM* cells. C. Silence of *CUL1* eliminated the downregulation of DNA-PKcs after 2 Gy X-irradiation induced by RPRM. D. Silence of *FBXW11* eliminated the downregulation of DNA-PKcs after 2 Gy X-irradiation induced by RPRM. E. Pretreatment of MLN4924 for 24 h, a potent and selective inhibitor of NAE, eliminated the downregulation of DNA-PKcs in H460-*RPRM* cells after 2 Gy X-irradiation. F. Effect of MLN4924 pretreatment combined with *FBXW11* silence on DNA-PKcs level in H460-*RPRM* cells after 2 Gy X-irradiation. G. *FBXW11* silence decreased the micronucleus formation in X-irradiated H460-*RPRM* cells. H. Comparison of the growth of H460-NC and H460-*RPRM* xenografts treated with combination of MLN4924 and radiotherapy (a single dose of 8 Gy X-irradiation). n=6. Different letters indicate statistically significant differences among groups (a-h for *P* < 0.001; A-B for *P* < 0.01). Significance was evaluated via a Type III ANOVA with Tukey’s post-hoc correction in R (v4.2.2). I. Representative TUNEL staining in sections from H460-NC and H460-*RPRM* xenografts treated with MLN4924 combined with radiotherapy and quantification of TUNEL fluorescence intensity. n=3. Scale bar, 200 μm. \**P* < 0.05, \*\**P* < 0.01, \*\*\**P* < 0.001; significance was determined by two-way ANOVA followed by Tukey’s test or paired Student’s t-test (I).

To determine whether CUL1 and FBXW11 participate in DNA-PKcs degradation promoted by RPRM, we silenced *CUL1* and *FBXW11* in *RPRM*-overexpressing cells (Figure S4A and S4B), and found that knocking down either *CUL1* or *FBXW11* eliminated the downregulation of DNA-PKcs induced by RPRM in irradiated cells (Figure 3C, 3D and S4C). We also treated cells with MLN4924 prior to irradiation, a potent and selective inhibitor of NEDD8-activating enzyme (NAE) that is essential for its conjugation with CUL1 [42], and observed that DNA-PKcs level was restored in irradiated H460-*RPRM* cells after MLN4924 treatment (Figure 3E). Moreover, combination of MLN4924 pretreatment and *FBXW11* knockdown did not exhibit a synergistic effect in restoring DNA-PKcs level after IR (Figure 3F). These results indicate that RPRM downregulates DNA-PKcs through SCF^FBXW11^-mediated degradation.

We further confirmed the involvement of SCF^FBXW11^ complex in RPRM-mediated radiation response. *In vitro*, knocking down *FBXW11* in H460-*RPRM* cells decreased the micronucleus formation of cells following irradiation (Figure 3G), confirming a critical role of FBXW11 in RPRM-regulated DNA damage repair and radiosensitivity via mediating DNA-PKcs degradation. *In vivo*, we evaluated the effect of combining MLN4924 with radiotherapy using xenografts formed by H460-NC and H460-*RPRM* cells. As shown in Figure 3H, *RPRM* overexpression significantly suppressed tumor growth. Furthermore, radiotherapy induced stronger tumor growth inhibition in H460-*RPRM* xenografts than in H460-NC xenografts, consistent with our previous finding [21]. Importantly, while MLN4924 alone inhibited the growth of both H460-*RPRM* and H460-NC xenografts, the combination of MLN4924 and IR resulted in synergistic growth suppression only in H460-NC xenografts. Conversely, MLN4924 treatment significantly attenuated the radiosensitivity of H460-*RPRM* xenografts (Figure 3H and S5). In addition, in contrast to severe apoptosis in H460-NC xenografts, the combination of MLN4924 and IR only induced slight apoptosis in *RPRM*-overexpressing xenografts (Figure 3I). These results indicate that the efficacy of MLN4924 combined with radiotherapy is associated with the *RPRM* expression level in cancer cells. These findings further support our model wherein RPRM inhibits the NHEJ repair pathway through promoting the SCF^FBXW11^ ligase complex-mediated DNA-PKcs degradation.

### FBXW11 recognizes p-DNA-PKcs (T2609) and mediates its degradation

To confirm DNA-PKcs as a degradation substrate of FBXW11, we began by assessing whether an interaction occurred between FBXW11 and DNA-PKcs. Not unexpectedly, FBXW11 did bind to DNA-PKcs (Figure 4A and S6A). Furthermore, it has been reported that FBXW11 recognizes and binds to phosphorylated target proteins such as phosphorylated CTNNB1, NFKB1, DEPTOR, TFE3, MITF, etc [43–47]. DNA-PKcs has two clusters of phosphorylation sites, i.e., PQR around the S2056 site and ABCDE around the T2609 site. Phosphorylation of the two clusters is critical for the role DNA-PKcs plays in chromosomal end-processing and end-ligation [48]. Thus, we examined whether FBXW11 interacts with DNA-PKcs phosphorylated at T2609 or S2056. As shown in Figure 4B, 4C and S6B, FBXW11 did bind to p-DNA-PKcs (T2609) but not p-DNA-PKcs (S2056). Moreover, the interaction between FBXW11 and p-DNA-PKcs (T2609) was enhanced significantly in H460-*RPRM* cells post X-irradiation. This indicates that FBXW11 recognizes DNA-PKcs phosphorylated at T2609 but not at S2056. Accordingly, following irradiation, the phosphorylation level of DNA-PKcs at S2056 in H460-*RPRM* cells remained unchanged relative to that in H460-NC cells (Figure 4D). In contrast, the phosphorylation level of DNA-PKcs at T2609 in irradiated H460-*RPRM* cells was significantly decreased compared to that in irradiated H460-NC cells (Figure 4E, 4F). Furthermore, p-DNA-PKcs (T2609) ubiquitylation was significantly increased in H460-*RPRM* cells but not in H460-NC cells following irradiation (Figure 4G). Consistently, compared to their NC cells, A549-*RPRM* and AGS-*RPRM* cells also exhibited a significant reduction in phosphorylation at the T2609, but not the S2609, site of DNA-PKcs after IR (Figure S7). In addition, pretreatment with NU7441 prior to irradiation, a selective inhibitor of DNA-PKcs, dramatically decreased p-DNA-PKcs (T2609) levels in both irradiated H460-RPRM and NC cells, meanwhile eliminated the reduction of DNA-PKcs level induced by *RPRM* overexpression after IR (Figure 4H). These results indicate that FBXW11 recognizes p-DNA-PKcs (T2609) and mediates its ubiquitylation and degradation, leading to a reduction in the total level of DNA-PKcs after irradiation.

**Figure 4.**
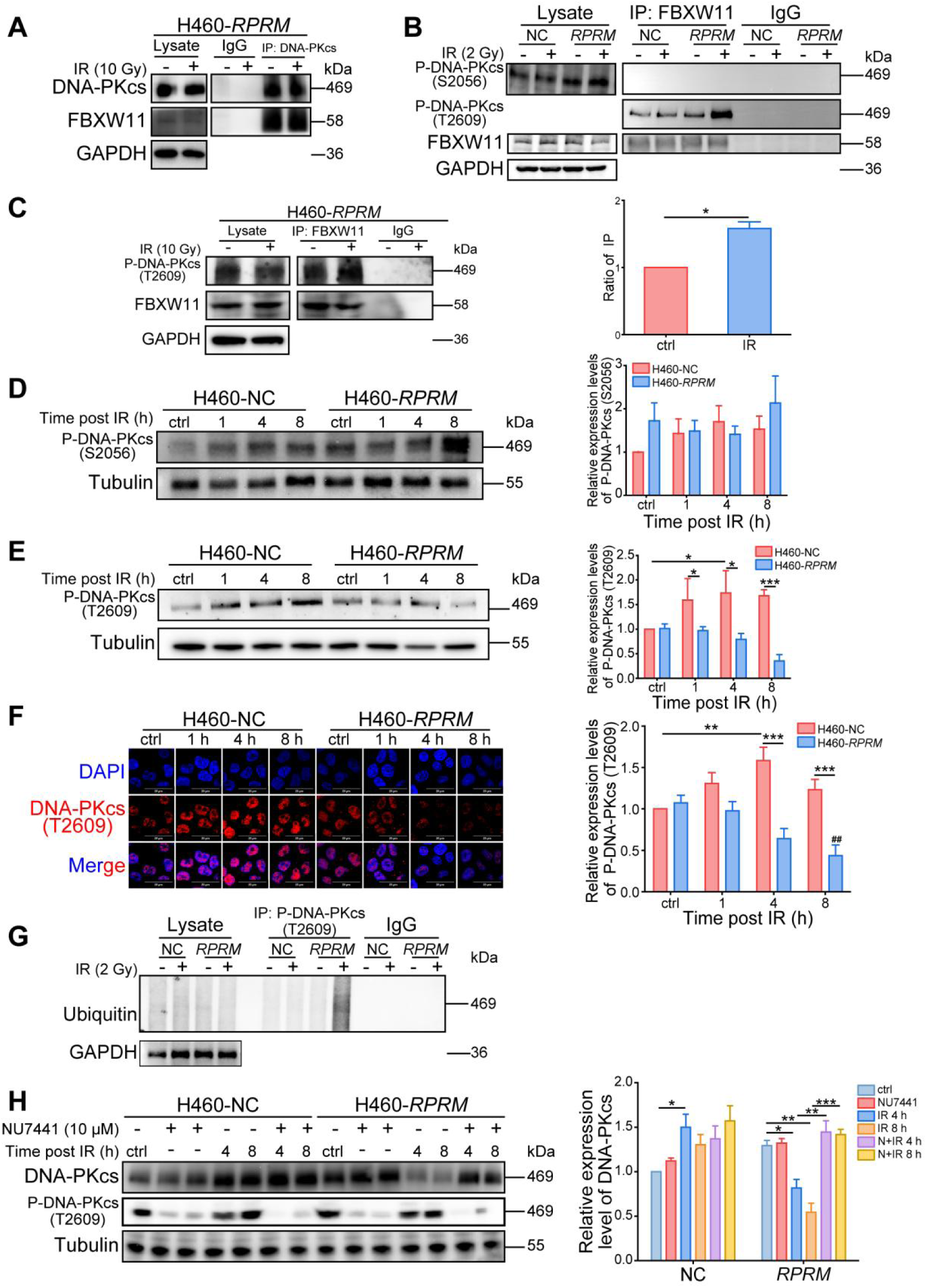
FBXW11 recognizes p-DNA-PKcs (T2609) and mediates its degradation. A. Co-IP demonstrating an interaction between DNA-PKcs and FBXW11. B. Co-IP demonstrating that FBXW11 bound to p-DNA-PKcs (T2609) but not p-DNA-PKcs (S2056). C. Interaction between FBXW11 and p-DNA-PKcs (T2609) was enhanced after IR. D. Phosphorylation levels of DNA-PKcs at S2056 in H460-NC/*RPRM* cells after IR determined by Western blot. E. Phosphorylation levels of DNA-PKcs at T2609 in H460-NC/*RPRM* cells after IR determined by Western blot. F. Representative immunofluorescence images of p-DNA-PKcs (T2609) in H460-NC/*RPRM* cells after IR and quantification of immunofluorescence intensity. Scale bar, 20 μm. Two hundred cells were analyzed for each group. G. Co-IP assessing p-DNA-PKcs (T2609) ubiquitylation in H460-NC/*RPRM* cells after IR. H. Effect of NU7441 on the levels of DNA-PKcs in H460-NC/*RPRM* cells after IR. \**P* < 0.05, \*\**P* < 0.01, \*\*\**P* < 0.001; ^##^*P* < 0.01, vs ctrl; significance was determined by two-way ANOVA followed by Tukey’s test.

### RPRM promotes the transcription of FBXW11 to enhance SCF^FBXW11^-mediated DNA-PKcs degradation

We then investigated how RPRM regulates SCF^FBXW11^ complex. Unexpectedly, RPRM did not affect the protein levels of both CUL1 (Figure 5A) and NEDD8 (Figure 5B). NEDD8-cullins complex level was slightly increased in H460-*RPRM* cells compared to H460-NC cells, yet the increase was not statistically significant (Figure 5C), indicating that RPRM did not obviously promote NEDD8-cullins conjugation before or after irradiation.

**Figure 5.**
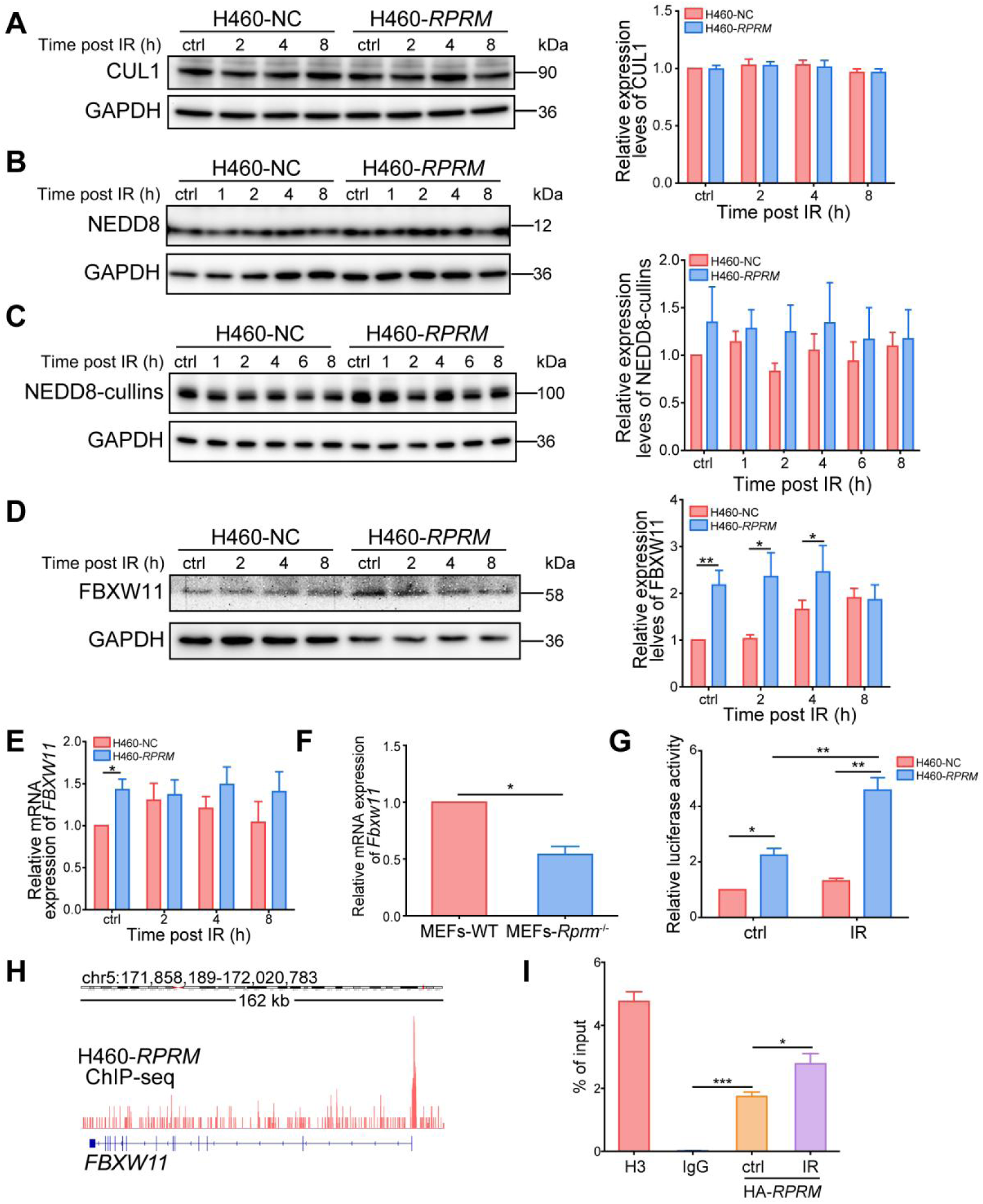
RPRM promotes *FBXW11* transcription but does not affect CUL1 protein. A. CUL1 protein levels in H460-NC/*RPRM* cells after 2 Gy X-irradiation. B. NEDD8 protein levels in H460-NC/*RPRM* cells after 2 Gy X-irradiation. C. The levels of NEDD8-cullins complex in H460-NC/*RPRM* cells after 2 Gy X-irradiation. D. FBXW11 protein levels in H460-NC/*RPRM* cells after 2 Gy X-irradiation. E. *FBXW11* mRNA levels in H460-NC/*RPRM* cells after 2 Gy X-irradiation. F. *Fbxw11* mRNA levels in MEFs-WT and MEFs-*Rprm*^-/-^. G. Luciferase reporter assay was performed to assess the transcription of the *FBXW11* promoter upon *RPRM* overexpression in H460 cells with/without IR. Data were normalized to Renilla luciferase activity. H. ChIP-seq on H460-*RPRM* cells reveals a specific binding peak for RPRM at the *FBXW11* promoter region. I. Independent ChIP-qPCR assay using non-irradiated and irradiated H460-*RPRM* cells (30 min post 2 Gy X-irradiation) confirms the enrichment observed by ChIP-seq. \**P* < 0.05, \*\**P* < 0.01, \*\*\**P* < 0.001; significance was determined by two-way ANOVA followed by Tukey’s test or paired t-test (F, I).

Instead, *RPRM* overexpression upregulated FBXW11 expression at both the mRNA and protein levels (Figure 5D, 5E). In contrast, knockout of *Rprm* gene significantly downregulated the transcription of *Fbxw11* in mouse embryonic fibroblasts (MEFs) (Figure 5F). To investigate whether RPRM protein regulates the transcription of *FBXW11* gene, we conducted a dual-luciferase reporter gene assay. The reporter plasmids containing the putative promoter sequence of *FBXW11* gene (*FBXW11*-Firefly luciferase reporter) and Renilla luciferase control (as an internal control) were co-transfected into both H460-NC and H460-*RPRM* cells. The groups co-transfected with NC-Firefly luciferase control vector and Renilla luciferase control vector served as the negative controls. As shown in Figure 5G, relative to the H460-NC control, *RPRM* overexpression led to a 2.2-fold (*P* < 0.05) and 4.6-fold (*P* < 0.01) enhancement in luciferase reporter activity before and after irradiation, respectively. Moreover, the combination of *RPRM* overexpression and IR led to a synergistic upregulation of *FBXW11* transcription. To further demonstrate that RPRM transcriptionally regulates *FBXW11*, we performed ChIP-seq and identified a significant peak enriched at the promoter of *FBXW11* gene (Figure 5H). The specific binding was further validated by independent qPCR assays, which also demonstrated its enhancement after X-irradiation (Figure 5I). These results confirm that RPRM specifically upregulates the transcription of *FBXW11*, which is enhanced by irradiation due to IR-induced RPRM nuclear translocation [21].

### RPRM nuclear translocation is essential for its negative regulation on DNA-PKcs

We have previously demonstrated that RPRM promotes ATM degradation following IR, in which RPRM nuclear import mediated by its phosphorylation status at serine 98 and importin transport receptor IPO11 is essential [21]. To clarify whether RPRM nuclear import is also required for its downregulation of DNA-PKcs, we employed the active (S98E) and inactive (S98A) RPRM point mutants. Compared to wild-type (WT) RPRM, the S98E mutant exhibited significantly reduced cytoplasm-to-nucleus translocation after irradiation, while the S98A mutant showed enhanced translocation (Figure S8). Consequently, in contrast to the RPRM-S98A mutant and WT RPRM, both of which downregulated DNA-PKcs after IR, the RPRM-S98E mutant failed to do so (Figure 6A). Similarly, in H460-*RPRM* cells, knockout of *IPO11*, which blocked RPRM nuclear import following IR (Figure S9) [21], dramatically increased the levels of DNA-PKcs relative to NC 4 and 8 h post irradiation, eliminating the negative regulatory effect of RPRM on DNA-PKcs (Figure 6B). These results demonstrate that RPRM nuclear import is essential for its suppression of DNA-PKcs. As expected, the nuclear translocation of RPRM was a prerequisite for its ability to upregulate FBXW11. As shown in Figure 6C and 6D, *IPO11* knockout significantly downregulated FBXW11 expression in *RPRM*-overexpressing cells at both the mRNA and protein levels. Subsequently, FBXW11-DNA-PKcs interaction was substantially inhibited in *IPO11-*KO cells (Figure 6E), indicating that RPRM nuclear import enhanced the interaction. These findings explain why inhibiting the nuclear import of RPRM restored DNA-PKcs levels (Figure 6B).

**Figure 6.**
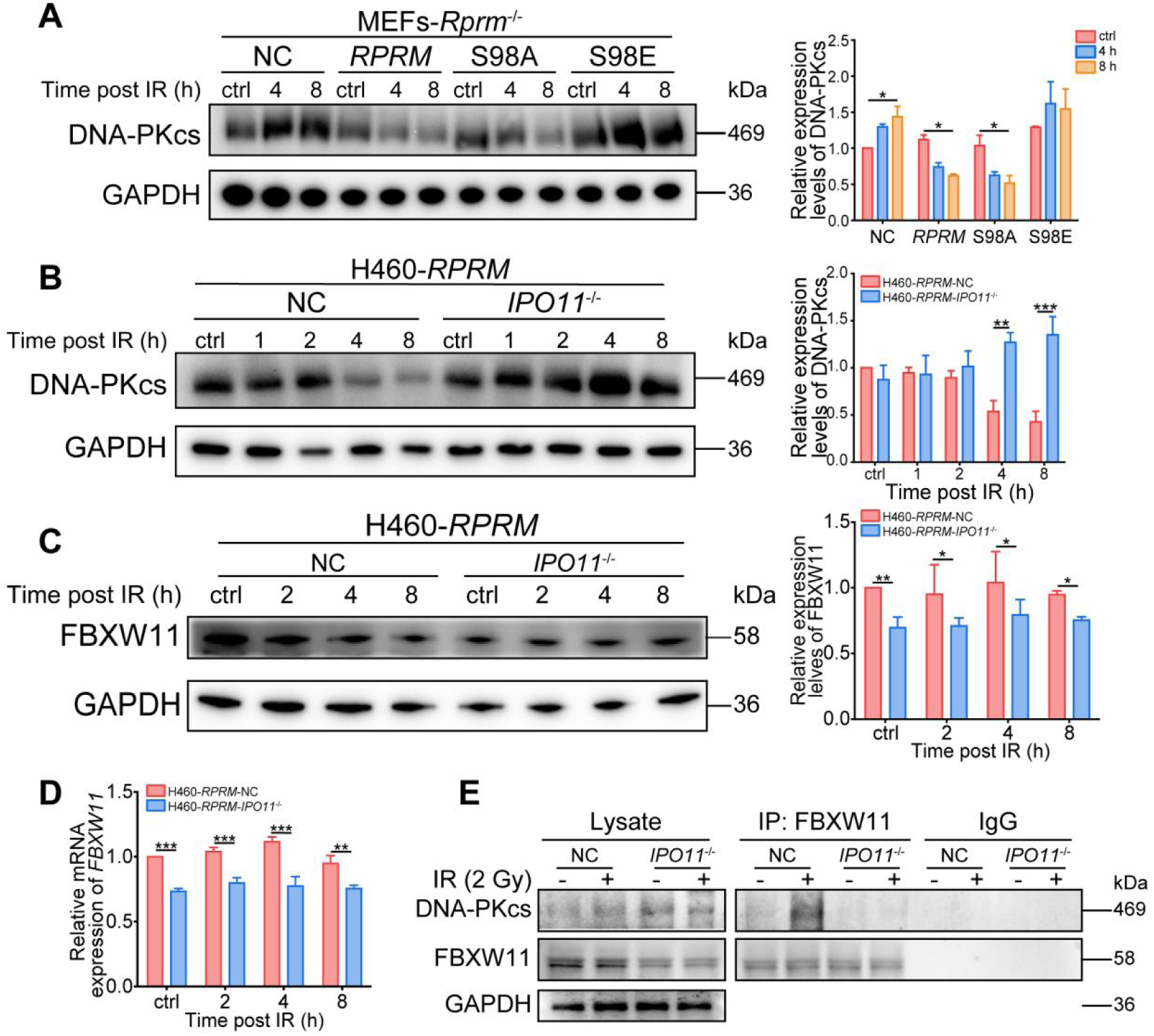
RPRM nuclear-translocation is required for DNA-PKcs inhibition. A. DNA-PKcs levels in MEFs-*Rprm*^-/-^ cells transfected with *RPRM*/S98A/S98E plasmids after 20 Gy X-irradiation. B. DNA-PKcs levels in H460-*RPRM*-NC/*IPO11*^-/-^ cells after 2 Gy X-irradiation. C. FBXW11 levels in H460-*RPRM*-NC/*IPO11*^-/-^ cells after 2 Gy X-irradiation. D. *FBXW11* mRNA levels in H460-*RPRM*-NC/*IPO11*^-/-^ cells after 2 Gy X-irradiation. E. Co-IP demonstrating that the interaction between FBXW11 and DNA-PKcs was eliminated in *IPO11*-knockout H460-*RPRM* cells after IR. \**P* < 0.05, \*\**P* < 0.01, \*\*\**P* < 0.001; significance was determined by two-way ANOVA followed by Tukey’s test.

### RPRM may be a biomarker for radiotherapy response in NSCLC

We uncovered that RPRM promotes the transcription of *FBXW11* and *PRKDC* in cancer cells. To validate whether this regulation holds true in cancer tissues, we analyzed the mRNA expression of *RPRM*, *FBXW11* and *PRKDC* in 20 NSCLC patient samples to evaluate the correlation between their levels. As shown in Figure 7A–7C, *RPRM* expression demonstrated a significant positive correlation with both *FBXW11* expression (Spearman’s rho = 0.7965, *P* < 0.0001) and *PRKDC* expression (Spearman’s rho = 0.6338, *P* < 0.01) in the NSCLC patient cohort. These findings validate our observations in cancer cells *in vitro* (Figure 2A, 5E), confirming an important role of RPRM in regulating gene transcription.

**Figure 7.**
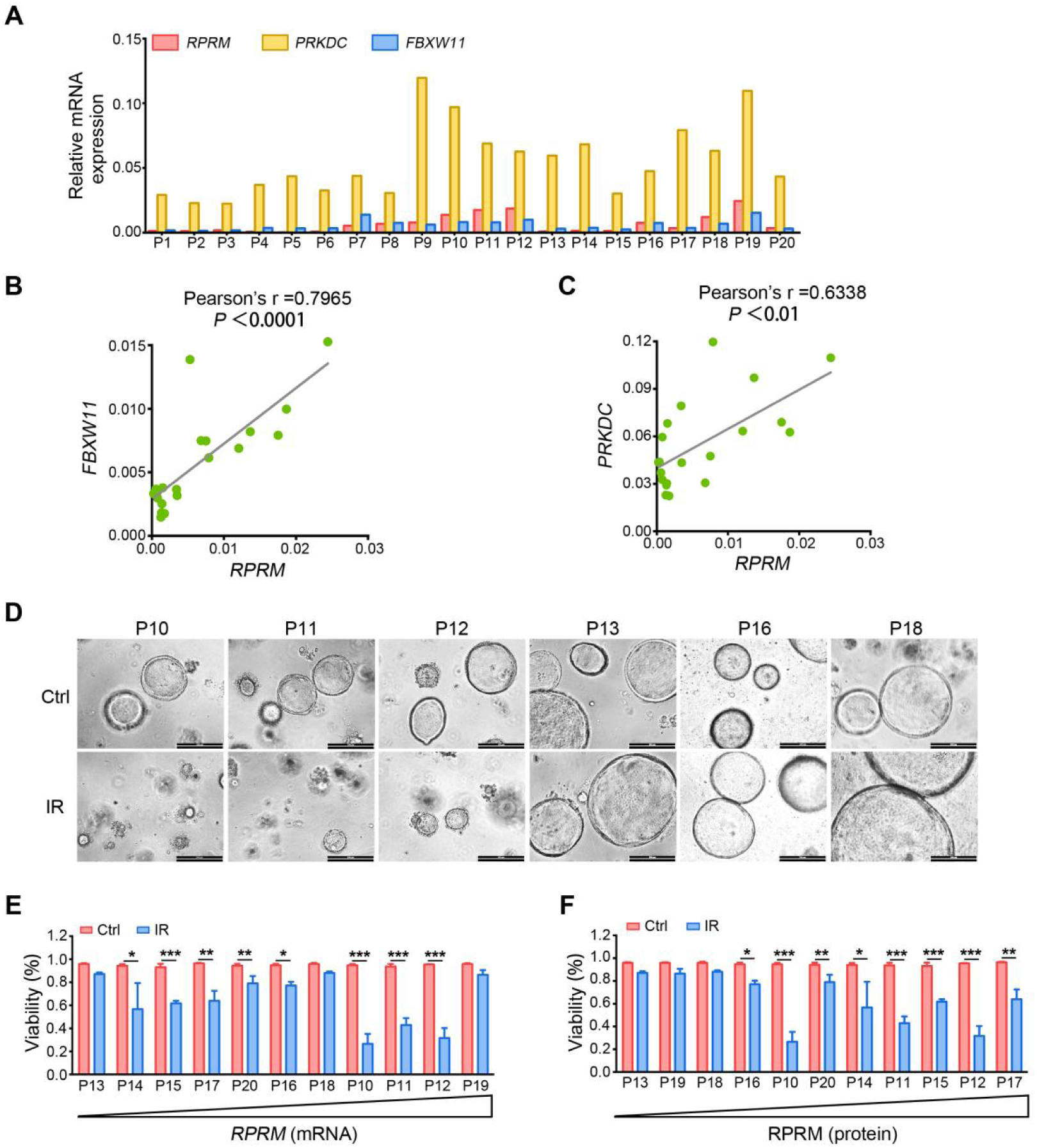
RPRM is related to PDO radiosensitivity. A. mRNA expression of *RPRM*/*FBXW11*/*PRKDC* detected in tumors from NSCLC patients. B. Correlation between the mRNA expression of *RPRM* and *FBXW11*. C. Correlation between the mRNA expression of *RPRM* and *PRKDC*. D. Eleven NSCLC PDOs were irradiated (0 and 5 Gy) and monitored for 14 days. Representative brightfield images of PDOs before and 14 days post irradiation. Scale bars, 200 μm. E. Correlation between PDO viability and *RPRM* mRNA expression. F. Correlation between PDO viability and RPRM protein expression. \**P* <0.05, \*\**P* <0.01, and \*\*\**P* <0.001; significance was determined by unpaired Student’s t-test.

To further investigate the impact of RPRM on cancer radiation response, we established patient-derived organoids (PDOs) and irradiated them with X-rays. As shown in Figure 7D–7F, the radiosensitivity of PDOs was more closely associated with RPRM protein levels than with its mRNA levels. PDOs with higher RPRM protein level showed more significant radiation response. This finding is consistent with RPRM’s role as a negative regulator of both DNA-PKcs (Figure 2) and ATM [21]. These results implicate RPRM as both a biomarker for evaluating radiation response and a therapeutic target in cancer radiotherapy.

## Discussion

In this study, we revealed an innovative mechanism in which protein RPRM suppresses the NHEJ repair pathway through its negative regulation of DNA-PKcs. Despite its designation as a tumor suppressor regulating key cellular processes such as cell cycle arrest, proliferation, cell death, migration, DDR, etc [21–34], RPRM’s core nature, molecular mechanisms and regulatory pathways are largely undefined. Here, we demonstrated that RPRM potently inhibits NHEJ by promoting proteasomal degradation of phosphorylated DNA-PKcs (T2609) mediated by SCF^FBXW11^ E3 ligase complex. These results extend RPRM’s established negative regulation of ATM [21] to DNA-PKcs, thus, emphasizing that RPRM, a poorly-characterized protein, functions as a key upstream regulator of PIKKs and plays a pivotal role in DNA damage repair, although whether RPRM impacts ATR remains to be investigated. Notably, the tumor organoids derived from cancer patients exhibited positive correlation between radiosensitivity and RPRM expression level, which is consistent with our previous finding that *RPRM* overexpression sensitizes cells to IR whereas *RPRM* (*Rprm*) deficiency induces radioresistance [21], indicating a potential implication of RPRM in cancer radiotherapy.

Along with our previous studies [21,34], we have revealed that RPRM regulates proteasomal degradation of proteins including ATM [21], CREB [34] and DNA-PKcs. Interestingly, RPRM directly binds to both ATM and CREB [21,34], but not to DNA-PKcs, suggesting that it mediates the degradation of these proteins through distinct pathways. The molecular mechanisms for ATM and CREB degradation promoted by RPRM remain elucidated. However, here we revealed that RPRM promotes DNA-PKcs degradation specifically through the SCF^FBXW11^ complex-mediated pathway.

Extensive research has revealed the key roles of DNA-PKcs in DNA damage repair, transcriptional regulation, DNA fidelity, metabolic regulation, cell death, adaptive immunity, and tumor microenvironment [49]. Nevertheless, the regulation of this critical multifunctional protein is poorly investigated so far. Feng et al. recently reported degradation of DNA-PKcs by the CUL4A-DTL ligase, impairing the NHEJ repair [18]. Our present finding identified the SCF^FBXW11^ ligase as an additional E3 ubiquitin ligase responsible for DNA-PKcs degradation. This indicates that DNA-PKcs can be degraded by different cullin-RING-ligase family members [50], utilizing their respective substrate receptor DTL [18] and F-box protein FBXW11. While DTL may recognize DNA-PKcs loaded on DNA [18,51], FBXW11 targets phosphorylated DNA-PKcs at T2609. As the substrate recognition component of SCF E3 ligase complex, FBXW11 recognizes phosphorylated target proteins such as TFE3 and MITF-A [52], and DEPTOR [45,53]. Thus, as expected, FBXW11 recognized p-DNA-PKcs (T2609), leading to its ubiquitylation and degradation, thereby reducing total DNA-PKcs levels. Interestingly, our findings also demonstrated that FBXW11 failed to recognize DNA-PKcs phosphorylated at S2056; consequently, we did not observe a reduction in p-DNA-PKcs (S2056) levels. The mechanistic basis for the selective recognition of p-DNA-PKcs (T2609) but not p-DNA-PKcs (S2056) by FBXW11 remains unclear and warrants further investigation. Given that phosphorylation of the T2609 cluster enhances DNA end processing, whereas phosphorylation at the S2056 cluster inhibits this activity [54], our results may suggest that the SCF^FBXW11^ ligase complex suppresses DNA end joining by targeting p-DNA-PKcs (T2609) for degradation, thereby explaining how RPRM inhibits the NHEJ repair pathway.

Consistent with previous study [39], we demonstrated that RPRM interacts with FBXW11 and CUL1. This would facilitate the assembly and stability of SCF^FBXW11^ complex. Moreover, RPRM promoted its transcriptional upregulation, resulting in increased FBXW11 protein levels, which promoted the degradation of p-DNA-PKcs (T2609). Although motif analysis following ChIP-seq indicate that RPRM does not appear to be a sequence-specific transcription factor (data not shown), its positive regulation of *FBXW11* transcription, as evidenced by dual-luciferase reporter and ChIP-qPCR assays, leads us to propose that RPRM may function as a transcriptional enhancer ‒ a novel function previously unreported. The fact that *RPRM* overexpression increased *PRKDC* expression provides another example, although further investigation is needed. Therefore, our results reveal key characteristics of RPRM protein.

In contrast to FBXW11, despite the interaction between RPRM and CUL1, *RPRM* overexpression did not alter the expression and activity of CUL1. As an inhibitor of NEDD8-activating enzyme (NAE) for Cullin proteins including CUL1, MLN4924 significantly inhibits tumor growth [55,56]. Our findings further demonstrated its therapeutic potential in NSCLC cell model, regardless of *RPRM* expression. However, when combined with IR, MLN4924 exhibited divergent effect: it radiosensitized control (NC) tumors but induced radioresistance in *RPRM*-overexpressing tumors. This paradoxical effect may be explained by RPRM’s inhibition of the NHEJ repair pathway, mediated through its promotion of the degradation of phosphorylated DNA-PKcs (T2609) following radiation exposure, which requires CUL1. Coincidentally, MLN4924 has demonstrated opposing effects in combination with radiation: it acted as a radiosensitizer in prostate cancer cells yet conferred radioresistance in nasopharyngeal carcinoma (NPC) cells [57,58]. Notably, *RPRM* is frequently hypermethylated and consequently downregulated in prostate cancer [59]. Collectively, these findings suggest that *RPRM* expression level is a key determinant of whether MLN4924 enhances or diminishes the efficacy of radiotherapy, thus, *RPRM* expression may serve as a predictive biomarker for the therapeutic outcome of combining MLN4924 with radiotherapy.

Clinically, the expression of *RPRM* mRNA was found to be highly heterogeneous among cancer patients, as evidenced by TCGA data spanning several cancer types (e.g., NSCLC, breast and pancreatic cancer) and further validated in independent NSCLC patient samples. Most importantly, high *RPRM* expression predicts better prognosis in these cancer patients, indicating its crucial role in cancer therapy. Furthermore, the radiation sensitivity of patient-derived cancer organoids was associated with the levels of RPRM protein. This aligns with our previous study of the TCGA lung squamous cell carcinoma dataset, demonstrating that high *RPRM* expression correlates with better prognosis in patients receiving radiotherapy [21]. These observations support the role of RPRM as a key inhibitor of DNA repair, suppressing both HR and NHEJ through its downregulation of ATM and DNA-PKcs. Therefore, given its central function in DNA damage repair, the heterogeneous expression of *RPRM* may has profound implication for patient stratification and treatment strategy.

In summary, we demonstrated that RPRM, a novel biomarker for cancer patient prognosis, is a more potent suppressor of the NHEJ repair pathway than of the HR pathway. Furthermore, we elucidated the molecular mechanism by which it inhibits NHEJ. Specifically, following irradiation, RPRM protein translocates from the cytoplasm into the nucleus via IPO11, amplifies *FBXW11* transcription, and interacts with both CUL1 and FBXW11 proteins to scaffold the assembly of the SCF^FBXW11^ E3 ligase complex, enhances the recognition of DNA-PKcs phosphorylated at T2609 by FBXW11, subsequently promotes SCF^FBXW11^-mediated degradation of p-DNA-PKcs (T2609), resulting in reduced DNA-PKcs levels and impaired NHEJ repair (Figure 8). In addition, we validated that high RPRM level correlates with significant radiosensitivity in patient-derived cancer organoids. These data provided not only more insight into the characteristics and functions of RPRM but also a strong rationale for exploring RPRM as a companion biomarker and the RPRM-SCF^FBXW11^-DNA-PKcs axis as a therapeutic target to guide personalized cancer treatment.

**Figure 8.**
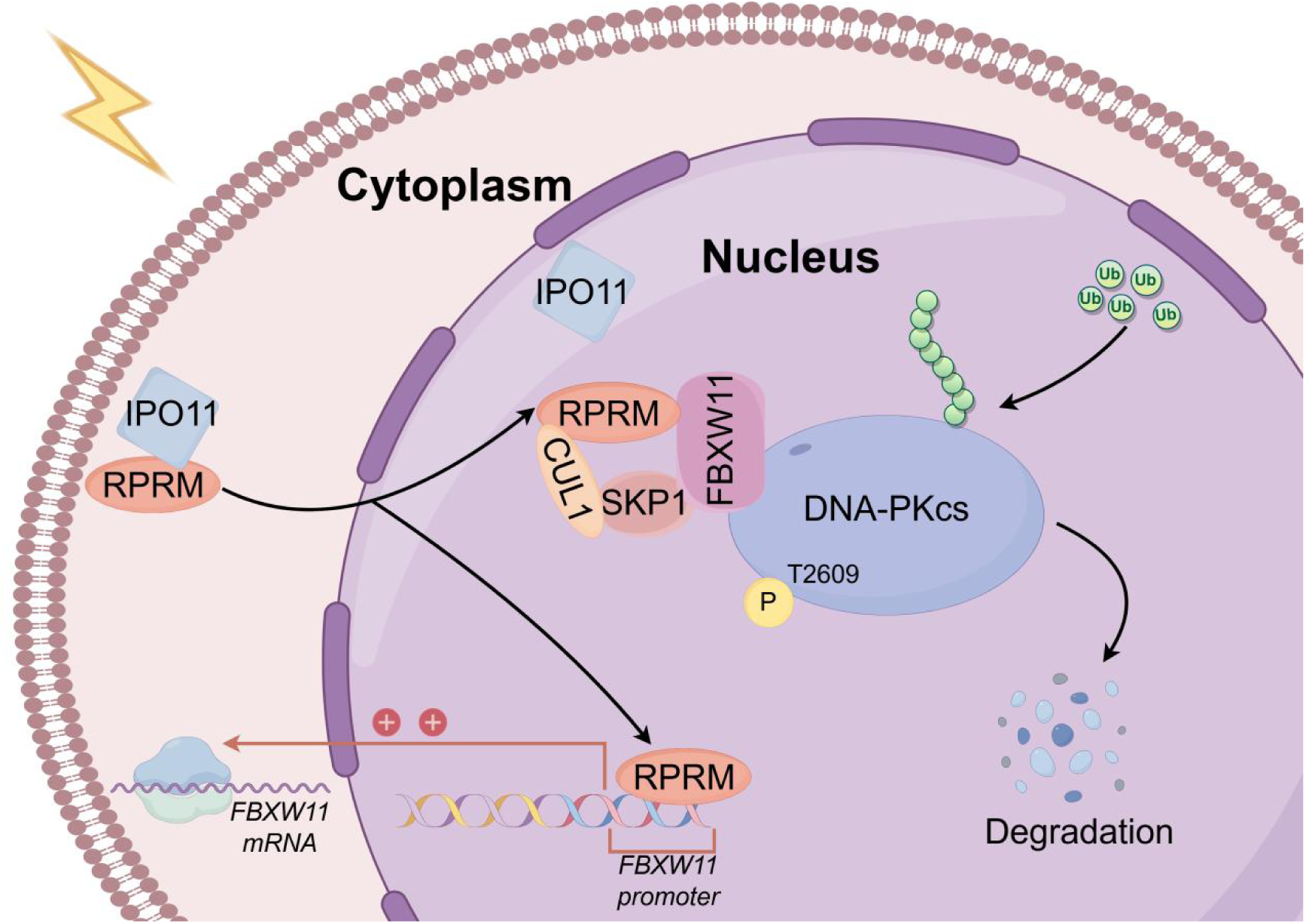
Schematic diagram of the mechanism by which RPRM promotes DNA-PKcs degradation through SCF^FBXW11^ complex.

### Limitations of the study

The present study reveals the novel mechanism by which RPRM suppresses the NHEJ repair pathway through promoting SCF^FBXW11^-mediated p-DNA-PKcs (T2609) degradation. We confirmed that RPRM interacts with both CUL1 and FBXW11, investigated how RPRM impacts CUL1 and FBXW11 as well as their roles in p-DNA-PKcs (T2609) degradation. However, future studies should delineate whether interaction between RPRM and CUL1 as well as FBXW11 is essential for the degradation of p-DNA-PKcs (T2609) promoted by RPRM. Another limitation of this study is that the precise molecular mechanism by which RPRM enhances the transcription of *FBXW11* remains to be fully elucidated. Although our luciferase reporter assay and ChIP data consistently demonstrate a transcriptional enhancement effect, the evidence suggests that RPRM does not appear to be a direct transcription factor. It may instead function as a co-factor or a scaffold that recruits other transcriptional activators to the promoter of *FBXW11*. Future work is needed to identify the direct binding partners of RPRM at this locus.

Moreover, it is important to note that the clinical correlation of RPRM with radiation response reported here is derived from NSCLC organoid models rather than direct patient data. While organoids faithfully recapitulate key tumor characteristics, they cannot fully mirror the complexities of the *in vivo* tumor microenvironment and systemic patient factors. Therefore, future investigations are necessary to extend these findings to patient cohorts and to correlate RPRM expression with clinical outcomes. Additionally, validation in larger, multi-center studies that encompass a spectrum of cancer types to confirm the broader applicability of RPRM as a biomarker and therapeutic target for radiotherapy is required.

## Methods

### Human specimens

The collection of tissue samples and all subsequent experiments were reviewed and approved by the Ethics Committee of Fujian Medical University and were conducted in accordance with obtained patient informed consent.

### Mouse experiments

All animal experiments were approved by the Ethics Committee of Soochow University and conducted in strict accordance with the University’s Medical Experimental Animal Care Guidelines, which adhere to China’s National Animal Welfare Policies. All mice were housed in the specific pathogen-free (SPF) facility at Soochow University’s Experimental Animal Center. Animals were maintained at 22 ± 2°C under a 12/12-hour light/dark cycle with food and water *ad libitum*. MEFs were isolated from C57BL/6 mouse embryos. Female BALB/c nude mice (4–5 weeks old) were used to establish subcutaneous tumor xenografts. H460-NC cells (3 × 10⁶) and H460-*RPRM* cells (5 × 10⁶) were injected subcutaneously into the right flanks of separate mouse groups. When tumors reached approximately 100 mm³, MLN4924 (0.12 mg in 100 μl per mouse) was administered via intratumoral injection. Twenty-four hours later, tumors were locally irradiated (8 Gy) with X-rays. Tumor dimensions (length and width) were then measured daily with digital calipers, and tumor volumes were calculated using the formula: Volume (mm³) = 1/2 × length × width². Mice were humanely euthanized when tumors reached ∼ 1000 mm³ or at 21 days post-irradiation (whichever occurred first), followed by tumor tissue harvest.

### Cell culture

All cell lines used in this study were confirmed mycoplasma-free and maintained in a humidified incubator at 37°C with 5% CO₂. Parental H460 cells were cultured in RPMI-1640 medium supplemented with 10% fetal bovine serum (FBS), 1% penicillin-streptomycin (PS), 10 mM HEPES, and 1 mM sodium pyruvate. H460-NC/*RPRM* cells were maintained in the same medium supplemented with 4 µg/ml puromycin. H460-*RPRM*-DRR (tet-on) cells were cultured in the same medium supplemented with 4 µg/ml puromycin and 200 µg/ml Geneticin (G418). H460-*RPRM*-NC/*IPO11*^-/-^ cells were maintained in the same medium supplemented with 4 µg/ml puromycin and 200 µg/ml hygromycin B.

AGS-NC/*RPRM* cells were maintained in F-12K medium supplemented with 10% FBS, 1% PS, and 2 µg/ml puromycin. A549-NC/*RPRM* cells were cultured in RPMI-1640 medium supplemented with 10% FBS, 1% PS, and 200 µg/ml hygromycin B.

As described previously [21], WT MEFs and *Rprm*^-/-^ MEFs were isolated from WT and *Rprm* KO mouse embryos and maintained in Dulbecco’s Modified Eagle Medium (DMEM, high glucose) supplemented with 10% FBS and 1% PS. All MEFs used in experiments were between passages 4 and 6.

To establish H460-*RPRM* (tet-on)-DRR cells, H460 cells were transduced with lentivirus LV-TetIIP-*RPRM* (NM_019845)-Puro. Transduced cells were selected in complete RPMI-1640 medium supplemented with 8 µg/ml puromycin. Single clone exhibiting strong doxycycline (DOX)-induced *RPRM* expression were selected and expanded into a cell line designated H460-*RPRM* (tet-on). H460-*RPRM* (tet-on) cells were then transfected with the plasmid pLCN DSB Repair Reporter (DRR). Transfected cells were selected in complete RPMI-1640 medium containing 400 µg/ml G418 and 8 µg/ml puromycin. A single clone was selected and expanded to establish the stable cell line H460-*RPRM* (tet-on)-DRR, which was then cultured in complete RPMI-1640 medium supplemented with 4 µg/ml puromycin and 200 µg/ml G418.

### Irradiation

Following irradiation with X-rays at designated doses using an RAD SOURCE RS2000 machine (160 kVp, dose rate: 1.225 Gy/min), cells or tumors were harvested at specified time points for subsequent analysis.

### Tumor tissue section preparation and TUNEL staining

Tumor tissues were fixed in 4% paraformaldehyde (PFA) at 4°C for 48 h, followed by sequential dehydration in a graded sucrose series (15% and 30%) with 24-hour immersions per step at 4°C. The dehydrated tissues were embedded in Optimal Cutting Temperature (OCT) compound, snap-frozen, and cryosectioned. The resulting sections were stored at-20°C for subsequent analysis.

For TUNEL staining, sections were first equilibrated to room temperature (RT) for 30 min. Apoptotic cells were labeled using a TUNEL Assay Kit (488 nm fluorescence), and nuclei were counterstained with DAPI. After mounting with an antifade medium, images were captured using an Olympus FV2000 laser scanning confocal microscope. The apoptosis level was quantified by measuring the relative fluorescence intensity of TUNEL-positive cells with ImageJ software.

### DSB repair reporter (DRR) assays for NHEJ and HR

H460-*RPRM* (tet-on)-DRR cells were seeded at a density targeting approximately 20-30% confluency after overnight culture. Cells were treated with 0.3 µg/ml DOX for 24 h to induce *RPRM* overexpression. Following DOX removal and washing, cells were co-transfected with the pCBASceI plasmid and HR donor at a 1:1 mass ratio using Lipo8000™. Twenty-four hours post-transfection, cells were trypsinized and resuspended in PBS. Cell suspensions were transferred to black opaque 96-well microplates (3 × 10⁴ cells per well). Green fluorescent protein (GFP) and mCherry fluorescence intensities were measured using a Synergy 2 microplate reader (Biotek) at appropriate excitation/emission wavelengths (488/530 nm for GFP; 580/620 nm for mCherry).

### Protein extraction, Co-Immunoprecipitation (Co-IP) and Western blotting (WB)

Cells were lysed on ice for 1 h with RIPA buffer supplemented with a protease inhibitor cocktail and PMSF (phosphatase inhibitors were included when analyzing phosphorylated proteins). Following centrifugation at 16,000 × g for 15 min at 4°C, the supernatant was collected. Protein concentration was determined using the Enhanced BCA Protein Assay Kit.

For Co-IP, 200–1000 μg of total protein was incubated overnight at 4℃ with the appropriate primary antibody or control IgG. The complexes were subsequently captured by incubation with Protein A/G Agarose beads for 3 h at 4°C under gentle agitation. After five washes with ice-cold RIPA buffer, the immunoprecipitated proteins were eluted and analyzed by Western blotting.

For Western blotting, equal amounts of protein (typically 20–50 µg) were denatured in 4× Loading buffer and resolved by 6% or 10% SDS-PAGE according to the molecular weight of the targeted proteins. The separated proteins were transferred onto polyvinylidene difluoride (PVDF) membranes. After blocking with 5% skim milk or 5% bovine serum albumin (BSA) in Tris-buffered saline containing 0.1% Tween-20 (TBST) for 2 h at RT, the membranes were incubated with primary antibodies diluted in antibody diluent overnight at 4°C, followed by probe with appropriate horseradish peroxidase (HRP)-conjugated secondary antibodies (diluted in blocking buffer) for 1 h at RT. Protein signals were detected using the FluorChem^TM^ M System (Alpha) with enhanced chemiluminescence (ECL) substrate and analyzed with ImageJ (NIH).

### Immunofluorescence (IF)

After treatment, cells on coverslips were fixed with 4% PFA for 15 min at RT. The cells were then permeabilized with PBS containing 1% Triton X-100 for 20 min at 4°C and blocked with 5% goat serum in PBS for 30 min at RT. The samples were incubated with primary antibodies for 1 h at RT, followed by incubation with corresponding fluorophore-conjugated secondary antibodies for 1 h at RT in the dark. Finally, coverslips were mounted on glass slides using an antifade mounting medium containing DAPI. Images were acquired using an Olympus FV2000 laser scanning confocal microscope.

### RNA extraction and Quantitative Real-Time PCR (qRT-PCR)

Total RNA was extracted from cells using TRIzol^®^ reagent. RNA concentration and purity were determined using the NanoDrop 2000c spectrophotometer (Thermo Fisher Scientific). Following the manufacturer’s instructions, complementary DNA (cDNA) was synthesized from 1 µg of the total RNA using the HiScript III All-in-one RT SuperMix Perfect for qPCR kit. Quantitative real-time PCR was performed using the Taq Pro Universal SYBR qPCR Master Mix on a real-time PCR detection system (ViiA™ 7 system, Thermo Fisher Scientific). Relative gene expression levels were calculated using the comparative Ct (ΔΔCt) method, with *GAPDH/Gapdh* or *ACTIN* serving as the internal reference genes. Gene-specific primer sequences used are listed in Table 4.

### Dual-luciferase reporter assay

Cells were co-transfected with either the NC-Firefly luciferase control or the *FBXW11*-Firefly luciferase reporter plasmid (*FBXW11*-promoter), along with a Renilla luciferase control plasmid for internal normalization. Fourty-eight hours post-transfection, firefly and renilla luciferase activities were measured sequentially using the Dual-Luciferase Reporter Assay Kit on a multifunctional microplate reader (synergy2, Biotek). Firefly luciferase activity was normalized to renilla luciferase activity for each sample. The relative luciferase activity of the *FBXW11* promoter was determined and presented as a bar graph, normalized to the ctrl-H460-NC-*FBXW11*-promoter group, which was set to 1.0.

### Chromatin immunoprecipitation (ChIP) followed by sequencing and quantitative PCR

Chromatin immunoprecipitation (ChIP) was performed using the BeyoChIP™ Enzymatic ChIP Assay Kit according to the manufacturer’s instructions. Briefly, there are several steps involved: cell culture and crosslinking, cell lysis and chromatin fragmentation, immunoprecipitation, elution and reverse crosslinking, and DNA purification. Immunoprecipitation was carried out using an anti-HA antibody, with an anti-Histone H3 antibody and normal IgG serving as positive and negative controls. The ChIP-enriched DNA libraries were constructed and sequenced by GENEWIZ (Suzhou) Biotechnology Co., Ltd. Based on the sequencing data, specific primers (listed in Table 4) were designed, and qPCR was performed to validate the enrichment of the *FBXW11* promoter region in the anti-HA ChIP samples.

### Micronucleus formation assay

Cells seeded on glass coverslips were irradiated. Twenty-four hours post-irradiation, the cells were fixed with freshly prepared fixative (methanol:acetic acid, 3:1) for 10 min at RT. The fixed cells were stained with DAPI for 2 min in the dark. Coverslips were mounted using an antifade mounting medium. Micronuclei were counted under a fluorescence microscope (Leica DM2000). Nuclei containing micronuclei were considered positive, and 1000 nuclei were counted per sample.

### Organoid culture and irradiation

Patient-derived NSCLC organoids were generated from resected tumor tissues. PDOs were cultures as previously described [60]. Briefly, tissue samples were minced and digested enzymatically. The resulting cell suspensions were kept on ice, resuspended in Matrigel at a 1:1 ratio, and plated as droplets in pre-warmed multi-well plates. The plates were briefly inverted to centralize the droplets, followed by incubation at 37°C to facilitate Matrigel polymerization. Upon solidification, the cultures were overlaid with complete growth medium. Organoids were exposed to X-rays (1.5 Gy/min, 160 kV, X-RAD 320) and the culture medium was replaced every three days after irradiation.

### Assessment of Organoid Viability

Following dissociation into single cells, organoids were seeded in 96-well plates at a density of 3,000 cells per well. Fouty-eight hours later, organoids were irradiated with 5 Gy X-rays. Organoid viability was determined 14 days after irradiation using the Organoid Vitality Assay Kit (HY-K6016, MCE, USA). After complete removal of the original medium, 100 μl of the organoid vitality working solution was added to each well, and the plates were incubated at 37°C for 2 hours. Wells containing only the dye solution without organoids were set as the negative control. Absorbance was then measured at 560 nm using a microplate reader. Viability was expressed as a percentage of the absorbance relative to that of non-irradiated controls, after subtracting the values of the negative controls. Data were derived from triplicate wells for each condition.

## Statistical Analysis

Data are presented as the mean ± SEM from at least three independent biological replicates. All statistical analyses and graphical presentations were generated using Origin 2021 software (OriginLab). For multiple-group comparisons, a two-way repeated-measures ANOVA with Tukey’s post hoc test was applied. Comparisons between two groups were performed using paired or unpaired Student’s t-test, as appropriate. A linear mixed-effects model was employed to analyze the longitudinal growth data of tumor xenografts. The model included treatment group, time, and their interaction as fixed effects, with mouse identity modeled as a random intercept. Significance was evaluated via a Type III ANOVA with Tukey’s post-hoc correction in R (v4.2.2). A *P-*value < 0.05 was considered statistically significant.

## Funding

This work was supported by the National Natural Science Foundation of China (Grant 32271279), Suzhou Bureau of Science and Technology (Suzhou Science and Technology Development Program Project, Jiangsu Finance-Education Document No. 160 (2023)), the Natural Science Foundation of Fujian Province (Grant 2024J01571)

Suzhou Fundamental Research Project (SJC2023001), Key Laboratory of Radiation Damage and Treatment of Jiangsu Provincial Universities and Colleges, and the Priority Academic Program Development of Jiangsu Higher Education Institution (PARD).

## Acknowledgements

We thank professors Jianjian Li and Shuyu Zhang for their valuable advice.

## Author contributions

Z.L. designed and performed the cellular and animal experiments. Y.Z. and B.T. analyzed the TCGA and NSCLC cohorts and conducted patient-derived organoid (PDO) experiments. S.X., Z.Z., and J.W. contributed to cellular and animal experiment design, data analysis and interpretation. G.O. and J.H. assisted in the TCGA analysis and PDO experiments. M.Q. designed and supervised the research involving clinical samples. H.Y. conceived and supervised the entire research project. All authors participated in preparing and editing the manuscript.

## Declaration of interests

The authors declare no competing interests except that H. Y., Y. Z., and J. W. hold a Chinese patent entitled “An establishment of RPRM knockout mouse model and its implications” (Patent No: ZL 2019 1 0405248.9).

## Notes

### Competing Interest Statement

The authors have declared no competing interest.

