## supplementary data for "RPRM (Reprimo) triggers SCF^FBXW11^-mediated DNA-PKcs degradation to block non-homologous end joining and radiosensitize tumors"

**Table S1 Clinical characteristics of the NSCLC patient cohort**

| Case | Sex | Age | Lung cancer type | Subtype | TNM |
|---|---|---|---|---|---|
| P1 | F | 60 | Invasive adenocarcinoma | Acinar | $T_{1b}N_0M_0$, Ⅰ$_{A2}$ |
| P2 | M | 49 | Invasive adenocarcinoma | Acinar30%+lepidic70% | $T_{1c}N_0M_0$, Ⅰ$_{A3}$ |
| P3 | F | 56 | Invasive adenocarcinoma | Acinar80%+lepidic20% | $T_{1a}N_0M_0$, Ⅰ$_{A1}$ |
| P4 | M | 60 | Invasive adenocarcinoma | Acinar50%+lepidic50% | $T_{1b}N_0M_0$, Ⅰ$_{A2}$ |
| P5 | F | 69 | Minimally invasive adenocarcinoma | | $T_{1b}N_0M_0$, Ⅰ$_{A2}$ |
| P6 | F | 68 | Invasive adenocarcinoma | Acinar40%+papillary35%+micro papillary20%+solid5% | $T_{1b}N_0M_0$, Ⅰ$_{A2}$ |
| P7 | M | 57 | Invasive adenocarcinoma | Solid45%+acinar30%+lepidic20%+micro papillary5% | $T_{1c}N_0M_0$, Ⅰ$_{A3}$ |
| P8 | F | 65 | Invasive adenocarcinoma | Acinar60%+lepidic20%+micro papillary20% | $T_{1b}N_0M_0$, Ⅰ$_{A2}$ |
| P9 | F | 60 | Invasive adenocarcinoma | Acinar60%+solid30%+micro papillary10% | $T_{1b}N_0M_0$, Ⅰ$_{A2}$ |
| P10 | F | 51 | Invasive adenocarcinoma | Acinar70%+lepidic20%+papillary5%+micro papillary5% | $T_{2a}N_0M_0$, Ⅰ$_{B}$ |
| P11 | F | 42 | Invasive adenocarcinoma | Acinar80%+papillary15%+complex acinar5% | $T_{1b}N_0M_0$, Ⅰ$_{A2}$ |
| P12 | F | 58 | Invasive adenocarcinoma | Acinar80%+lepidic20% | $T_{1b}N_0M_0$, Ⅰ$_{A2}$ |
| P13 | F | 48 | Invasive adenocarcinoma | Solid85%+complex glandular5%+acinar10% | $T_{1b}N_0M_0$, Ⅰ$_{A2}$ |
| P14 | F | 42 | Minimally invasive adenocarcinoma | | $T_{1a}N_0M_0$, Ⅰ$_{A1}$ |
| P15 | F | 54 | Invasive adenocarcinoma | Mucinous and non-mucinous | $T_{1c}N_0M_0$, Ⅰ$_{A3}$ |
| P16 | M | 60 | Invasive adenocarcinoma | Micro papillary40%+acinar30%+papillary20%+solid10% | $T_{2a}N_1M_0$, Ⅱ$_{B}$ |
| P17 | F | 46 | Invasive adenocarcinoma | Papillary40%+complex acinar40%+solid10%+micro papillary10% | $T_{1c}N_0M_0$, Ⅰ$_{A3}$ |
| P18 | F | 68 | Adenocarcinoma in situ | | $T_{1a}N_0M_0$, Ⅰ$_{A1}$ |
| P19 | M | 45 | Invasive adenocarcinoma | | $T_{1a}N_0M_0$, Ⅰ$_{A1}$ |
| P20 | F | 66 | Invasive adenocarcinoma | Acinar95%+micro papillary5% | $T_{1b}N_0M_0$, Ⅰ$_{A2}$ |

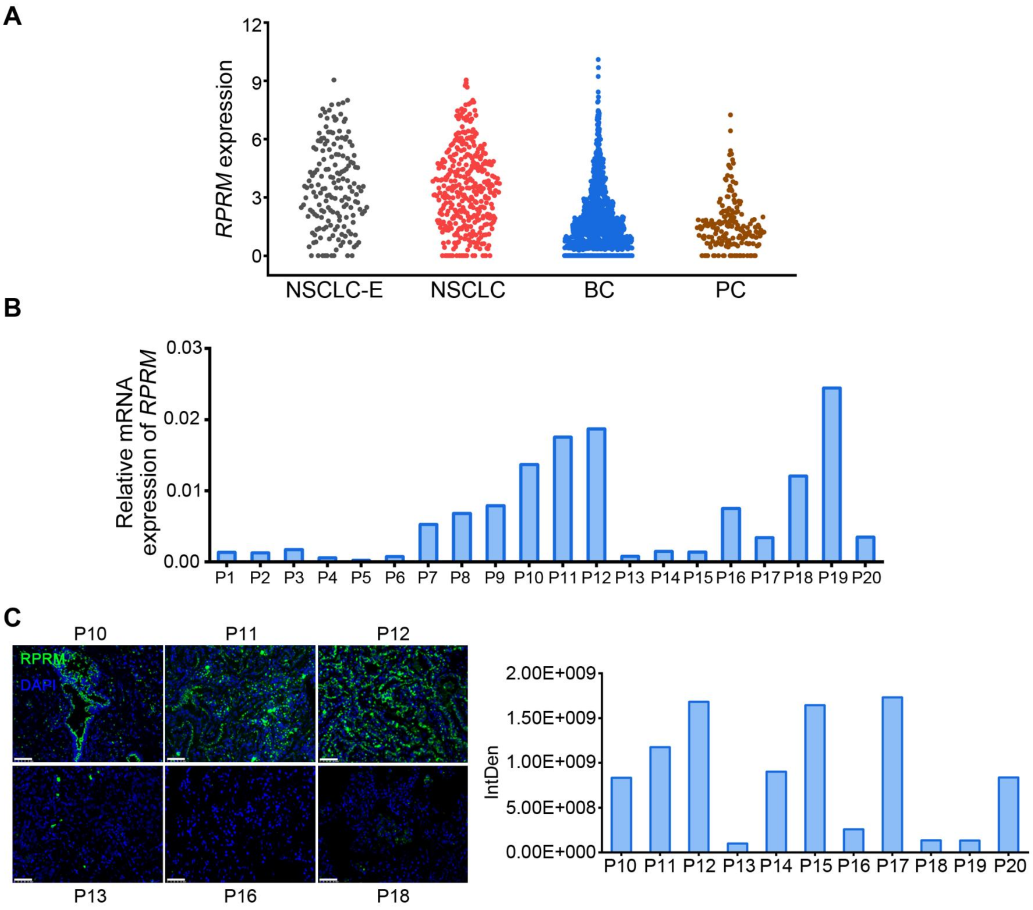


Figure S1. Inter-patient heterogeneity in *RPRM* gene expression across tumor tissues.

A. The TCGA dataset shows *RPRM* mRNA expression in various types of tumors including early stage of NSCLC (NSCLC-E), NSCLC, breast cancer (BC) and pancreatic cancer (PC).

B. *RPRM* mRNA expression detected in NSCLC samples (*Gapdh* was used as an internal control gene for normalization in the qRT-PCR).

C. Representative immunofluorescence images of RPRM in NSCLC samples and Pan-section quantification of RPRM fluorescence intensity using ImageJ. Scale bars, 50 μm.

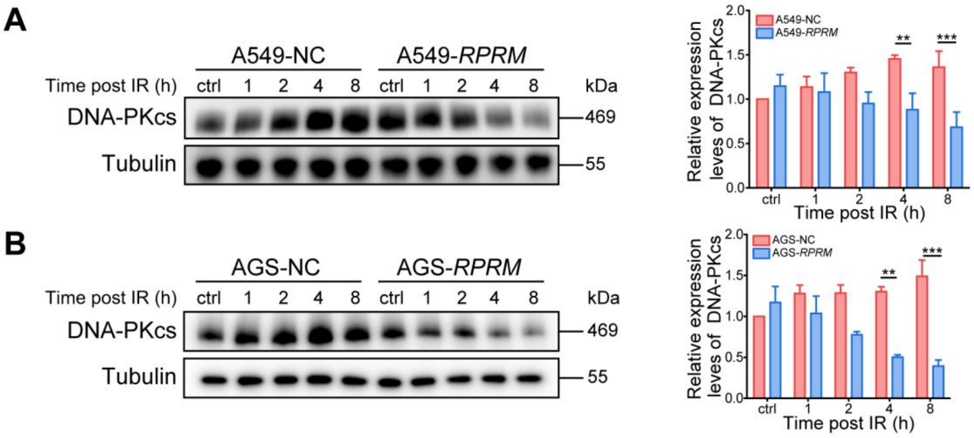


Figure S2. RPRM downregulates DNA-PKcs protein expression, related to Figure 2. The protein levels of DNA-PKcs in A549-NC/*RPRM* cells (A) and AGS-NC/*RPRM* cells (B) after 2 Gy X-irradiation. $^{**}P<0.01$, $^{***}P<0.001$; significance was determined by two-way ANOVA followed by Tukey’s test.

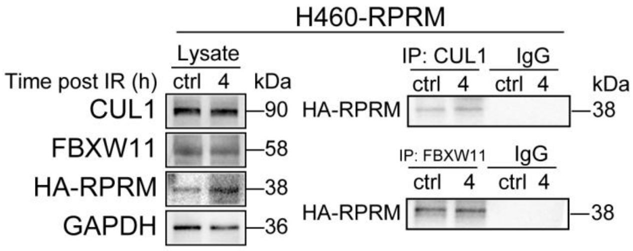


Figure S3. CUL1/FBXW11 binds to RPRM, related to Figure 3.

Co-IP was performed 4 h following 2 Gy X-irradiation to demonstrate the interaction between CUL1/FBXW11 and RPRM in H460-*RPRM* cells.

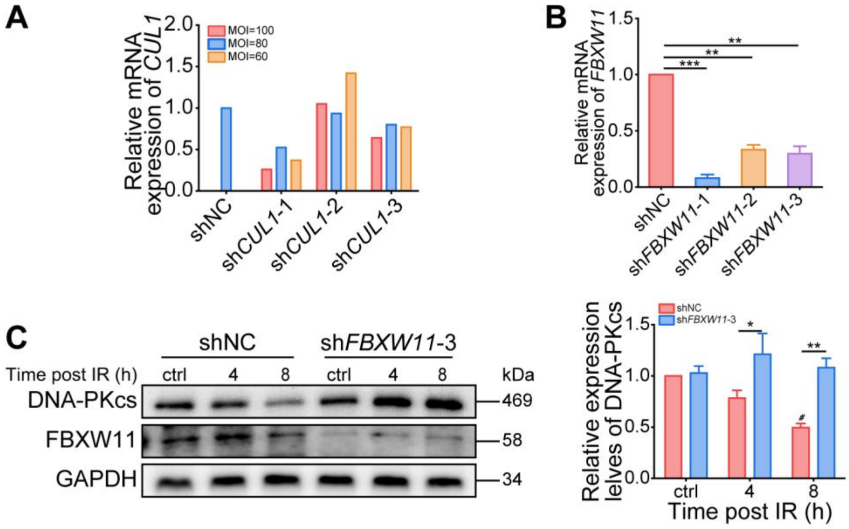


Figure S4. CUL1 and FBXW11 are essential for DNA-PKcs degradation, related to Figure 3.

A. The mRNA levels of *CUL1* in H460-*RPRM* cells after transduced with lentiviral vectors encoding distinct shRNA sequences against *CUL1* at various MOIs.

B. The mRNA levels of *FBXW11* in H460-*RPRM* cells after transduced with lentiviral vectors encoding distinct shRNA sequences against *FBXW11* at an MOI of 100.

C. The protein levels of DNA-PKcs in *FBXW11*-silenced H460-*RPRM* cells by #3 shRNA construct after 2 Gy X-irradiation.

*$P$<0.05, **$P$<0.01, ***$P$<0.001; #$P$<0.05 vs ctrl; significance was determined by two-way ANOVA followed by Tukey's test.

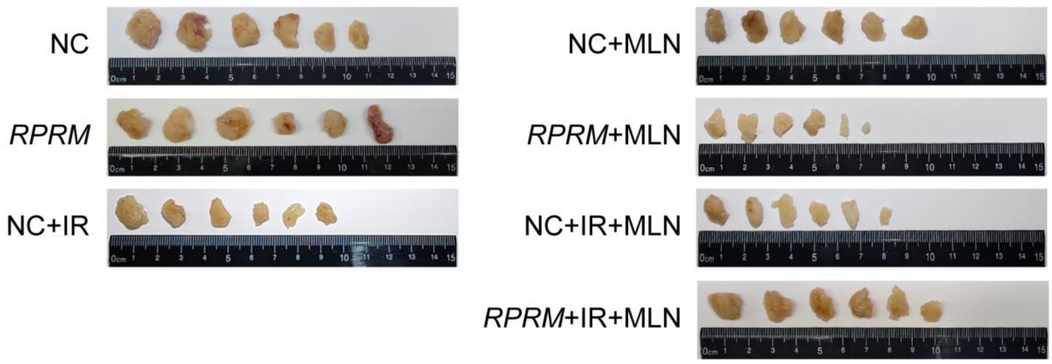


Figure S5. MLN4924 reduces the radiosensitivity of H460-*RPRM* xenografts, related to Figure 3.

The pictures of tumors excised on day 11 (NC), 14 (*RPRM*), 21 (all other groups) after IR, MLN4924 and combined treatment. Tumors of *RPRM*+IR group are not shown because they had completely regressed 17 days post-irradiation.

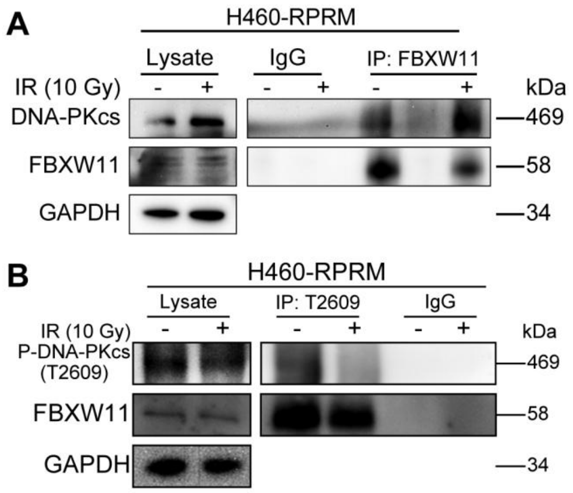


Figure S6. FBXW11 binds to DNA-PKcs/p-DNA-PKcs (T2609), related to Figure 4. Co-IP was performed 30 min after 10 Gy X-irradiation in H460-*RPRM* cells to demonstrate an interaction between FBXW11 and DNA-PKcs (A) as well as p-DNA-PKcs (T2609).

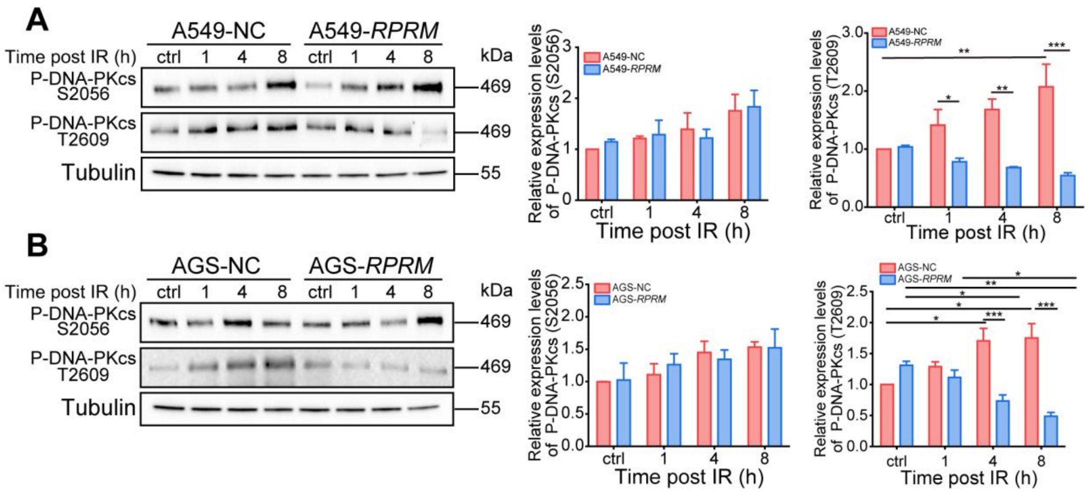


Figure S7. RPRM downregulates p-DNA-PKcs (T2609) but not p-DNA-PKcs (S2056) levels in irradiated cells, related to Figure 4.

The protein levels of p-DNA-PKcs (T2609/S2056) in A549-NC/*RPRM* cells (A) and AGS-NC/ *RPRM* cells (B) after 2 Gy X-irradiation. **P*<0.05, ***P*<0.01, ****P*<0.001; significance was determined by two-way ANOVA followed by Tukey's test.

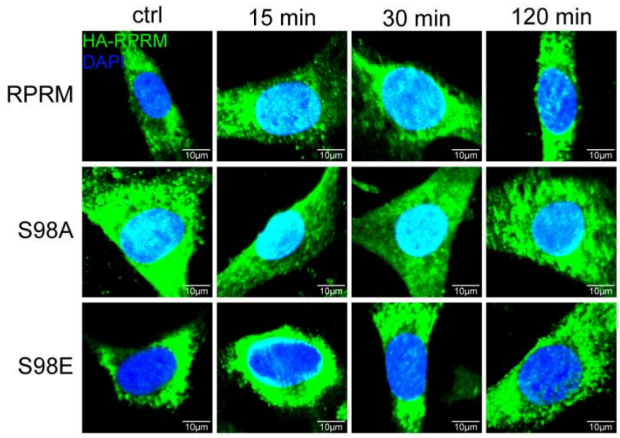


Figure S8. The phosphorylation status of RPRM affects its nuclear translocation following irradiation, related to Figure 6.

Representative immunofluorescence images of HA-RPRM in MEFs-*Rprm*$^{-/-}$ cells transfected with RPRM/S98A/S98E plasmids at different time points after 20 Gy X-irradiation. Scale bars, 10 μm

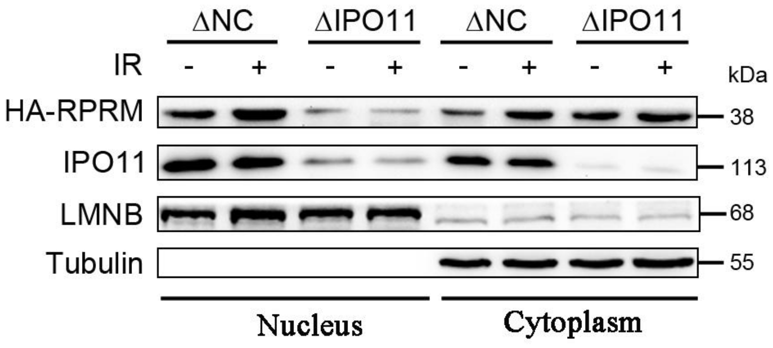


Figure S9. *IPO11* knockout blocks the nuclear import of RPRM induced by ionizing radiation, related to Figure 6.

The protein levels of HA-RPRM in the cytoplasm and nucleus of H460-*IPO11* deletion/negative control (ΔIPO11/ΔNC) cells 30 minutes after exposure to 2 Gy of X-ray irradiation.
