## supplementary materials for "RPRM (Reprimo) triggers SCF^FBXW11^-mediated DNA-PKcs degradation to block non-homologous end joining and radiosensitize tumors"

**Table 1**

Primary and secondary antibodies used for WB, Co-IP , ChIP and IF.

| Antibody Name | Catalog Number | Brand | Dilution/Usage | Host Species |
|---|---|---|---|---|
| DNA-PKcs | AF1888 | Beyotime | 1:1000(WB)<br>1:100 (IF) | Rabbit |
| α-Tubulin | 66031-1-Ig | Proteintech | 1:40,000 (WB) | Mouse |
| Ubiquitin | AF1705 | Beyotime | 1:1000 (WB) | Rabbit |
| Mouse IgG | A7028 | Beyotime | 0.5 μg (IP) | Mouse |
| Rabbit IgG | A7016 | Beyotime | 0.5 μg (IP) | Rabbit |
| GAPDH | AF1186 | Beyotime | 1:1000 (WB) | Mouse |
| DNA-PKcs | ab70250 | Abcam | 2 μg (IP) | Rabbit |
| HA-Tag | 66006-2-Ig | Proteintech | 1:800 (IF)<br>1 μg (IP) | Mouse |
| HA-Tag | AH158 | Beyotime | 1:50 (volume ratio, IP) | Mouse |
| Histone H3 | 81984-2-RR | Proteintech | 1 μg (IP) | Rabbit |
| FBXW11 | 13149-1-AP | Proteintech | 1:1000 (WB)<br>1 μg (IP) | Rabbit |
| FBXW11 | DF13009 | Affinity | 1：2000 (IF) | Rabbit |

| Antibody Name | Catalog Number | Brand | Dilution/Usage | Host Species |
|---|---|---|---|---|
| CUL1 | 12895-1-AP | Proteintech | 1:5000 (WB)<br>1 μg (IP) | Rabbit |
| NEDD8 | AF7557 | Beyotime | 1:1000 (WB) | Rabbit |
| NEDD8-cullins | ab81264 | Abcam | 1:2000 (WB) | Rabbit |
| Phospho-DNA-PKcs (T2609) | GTX24194 | GeneTex | 1:1000 (WB)<br>1:200 (IF)<br>1 μg (IP) | Rabbit |
| Phospho-DNA-PKcs (S2056) | GTX132793 | GeneTex | 1:1000 (WB) | Rabbit |
| RPRM | GTX110976 | GeneTex | 1:5000 (IF) | Rabbit |
| Goat-Anti-Mouse IgG (AlexaFluor 488) | RGAM002 | Proteintech | 1:500 (IF) | Goat |
| Goat-Anti-Rabbit IgG (AlexaFluor Cy3) | A0516 | Beyotime | 1:100 (IF) | Goat |
| Goat-Anti-Rabbit IgG-HRP | A0208 | Beyotime | 1:1000 (WB) | Goat |
| Goat-Anti-Mouse IgG-HRP | A0216 | Beyotime | 1:1000 (WB) | Goat |

**Table 2**

Animals in this study.

| Mouse Name | Source |
|---|---|
| BALB/c nude mice | Hangzhou Enlighten the Truth Laboratory Animal Technology Co., Ltd. |
| *Rprm*$^{-/-}$ C57BL/6 mice | Constructed in our previous study |

| Mouse Name | Source |
|---|---|
| BALB/c nude mice | Hangzhou Enlighten the Truth Laboratory Animal Technology Co., Ltd. |
| *Rprm*$^{-/-}$ C57BL/6 mice | Constructed in our previous study |
| WT C57BL/6 mice | Bred during the construction of KO mice in our previous study |

**Table 3**

Chemicals in this study.

| Reagent Name | Catalog Number | Source |
|---|---|---|
| MLN4924 | T6332 | TargetMol |
| Hygromycin B | 60224ES10 | Yeasen |
| Puromycin | ant-pr-1 | InvivoGen |
| G418 Sulfate | 60220ES03 | Yeasen |
| Doxorubicin (DOX) | E2516 | Selleck |
| Cycloheximide (CHX) | S7418 | Selleck |
| MG132 | S2619 | Selleck |
| NU7441 | HY-11006 | MedChemExpress |

**Table 4**

Primers in this study.

| Primer Name | Sequence (5'→3') |
|---|---|
| Human *RPRM*-Forward | CTGGCCCTGGGACAAAGAC |

| Primer Name | Sequence (5'→3') |
|---|---|
| Human *RPRM*-Reverse | TCAAAACGGTGTCACGGATGT |
| Human *ACTIN*-Forward | AAAAGCCACCCCACTTCTCTCT |
| Human *ACTIN*-Reverse | AATGCTATCACCTCCCCTGTGT |
| Human NEO-Forward | CGTTGGCTACCCGTGATATT |
| Human NEO-Reverse | GCCCAGTCATAGCCGAATAG |
| Human *CUL1*-Forward | CCACAGCAGCGGGTAGTATT |
| Human *CUL1*-Reverse | GCTCGAGCTTATCTGCGTGT |
| Human *FBXW11*-Forward | AGCTCATGTGTTCTGTGCCA |
| Human *FBXW11*-Reverse | ATTTGGAGGGCCATCTGTGG |
| Human *PRKDC*-Forward | GAACACCCTTTCCTGGTGAAG |
| Human *PRKDC*-Reverse | GGATCACTGGAGGTCATGGG |
| Mouse *Gapdh*-Forward | TGACCACAGTCCATGCCATC |
| Mouse *Gapdh*-Reverse | GACGGACACATTGGGGGTAG |
| Mouse *Fbxw11*-Forward | CCTGGTAGAGAGCATGTGCG |
| Mouse *Fbxw11*-Reverse | TGACGTTCCATTACTGATCTGG |
| Human *FBXW11*(promter)-Forward | GATCCGTGTCTTCCTGGTGG |
| Human *FBXW11*(promter)-Reverse | CACACAAGCGGAGACTCCTT |
| Human *FBXW11*(intron)-Forward | TGCTCTTTTTCACCCTCGCT |
| Human *FBXW11*(intron)-Reverse | CCCTCAAGCAGGTTCAGAGG |
| Human gene desert-Forward | ACATGTAGGCTGAGCACCAG |
| Human gene desert-Reverse | TCTTAGCACAGTGCCTGACG |

**Table 5**

Lentivirus in this study. All the following lentivirus are from Shanghai Genechem Co., Ltd.

| **Lentivirus Name** | **Top Strand Sequence (5'→3')** | **Bottom Strand Sequence (5'→3')** |
|---|---|---|
| LV-TetIIP-*RPRM*-Puro | ATGAATCCGGCCCTAGGCAACCAGACGGACGTGGCGGGCCTGTTCCTGGCCAACAGCAGCGAGGCGCTGGAGCGAGCCGTGCGCTGCTGCACCCAGGCGTCCGTGGTGACCGACGACGGCTTCGCGGAGGGAGGCCCGGACGAGCGTAGCCTGTACATAATGCGCGTGGTGCAGATCGCGGTCATGTGCGTGCTCTCACTCACCGTGGTCTTCGGCATCTTCTTCCTCGGCTGCAATCTGCTCATCAAGTCCGAGGGCATGATCAACTTCCTCGTGAAGGACCGGAGGCCGTCTAAGGAGGTGGAGGCGGTGGTCGTGGGGCCCTAC | |
| LV-NC-*FBXW11*-RNAi | GATCCGTTCTCCGAACGTGTCACGTAATTCAAGAGATTACGTGACACGTTCGGAGAATTTTTTC | AATTGAAAAAATTCTCCGAACGTGTCACGTAATCTCTTGAATTACGTGACACGTTCGGAGAACG |
| LV-*FBXW11*-RNAi-1 | CCGGTCGTACTCTCAATGGGCACAACTCGAGTTGTGCCCATTGAGAGTACGATTTTTG | AATTCAAAAATCGTACTCTCAATGGGCACAACTCGAGTTGTGCCCATTGAGAGTACGA |

| Lentivirus Name | **Top Strand Sequence (5'→3')** | **Bottom Strand Sequence (5'→3')** |
|---|---|---|
| LV-*FBXW11*-RNAi-2 | CCGGGTGTCATTGTAACTGGCTCTTCTCGAGAAGAGCCAGTTACAATGACACTTTTTG | AATTCAAAAAGTGTCATTGTAACTGGCTCTTCTCGAGAAGAGCCAGTTACAATGACAC |
| LV-*FBXW11*-RNAi-3 | CCGGGAACGAATGGTACGCACTGATCTCGAGATCAGTGCGTACCATTCGTTCTTTTTG | AATTCAAAAAGAACGAATGGTACGCACTGATCTCGAGATCAGTGCGTACCATTCGTTC |
| LV-NC-*CUL1*-RNAi | GATCCGTTCTCCGAACGTGTCACGTAATTCAAGAGATTACGTGACACGTTCGGAGAATTTTTTC | AATTGAAAAAATTCTCCGAACGTGTCACGTAATCTCTTGAATTACGTGACACGTTCGGAGAACG |
| LV-*CUL1*-RNAi-1 | CCGGGCCAGCATGATCTCCAAGTTACTCGAGTAACTTGGAGATCATGCTGGCTTTTTG | AATTCAAAAAGCCAGCATGATCTCCAAGTTACTCGAGTAACTTGGAGATCATGCTGGC |
| LV-*CUL1*-RNAi-2 | CCGGGCACACAAGATGAATTAGCAACTCGAGTTGCTAATTCATCTTGTGTGCTTTTTG | AATTCAAAAAGCACACAAGATGAATTAGCAACTCGAGTTGCTAATTCATCTTGTGTGC |
| LV-*CUL1*-RNAi-3 | CCGGGATTTGATGGATGAGAGTGTACTCGAGTACACTCTCATCCATCAAATCTTTTTG | AATTCAAAAAGATTTGATGGATGAGAGTGTACTCGAGTACACTCTCATCCATCAAATC |

**Table 6**

Plasmids in this study.

| Plasmid Name | Catalog Number | Source |
|---|---|---|
| pLCN DSB Repair Reporter (DRR) | #98895 | Addgene |

| Plasmid Name | Catalog Number | Source |
|---|---|---|
| pCAGGS DRR mCherry Donor EF1a BFP (HR Donor) | #98896 | Addgene |
| pCBASceI | #26477 | Addgene |
| NC (MEFs transfection control) | - | Constructed in our previous study |
| *RPRM* | - | Constructed in our previous study |
| *RPRM*-S98A mutant | - | Constructed in our previous study |
| *RPRM*-S98E mutant | - | Constructed in our previous study |
| NC-Firefly luciferase control | CON663 | Genechem |
| *FBXW11*-Firefly luciferase reporter (*FBXW11*-promoter) | K25E1404 | Genechem |
| Renilla luciferase control (RLUC) | CON696 | Genechem |

**Table 7**

Culture materials, reagents and kits in this study.

| Culture material/Reagent/Kit Name | Catalog Number | Source |
|---|---|---|
| Fetal bovine serum (FBS) | 164210-50 | Procell |
| RPMI-1640 medium | PM150110 | Procell |
| F-12K medium | PM150910 | Procell |
| DMEM medium | PM150210 | Procell |
| HEPES buffer | PB180325 | Procell |
| Sodium pyruvate | S8636 | Sigma |

| Culture material/Reagent/Kit Name | Catalog Number | Source |
|---|---|---|
| Penicillin-Streptomycin Solution (PS) | C0222 | Beyotime |
| Phosphate-buffered saline (PBS) | PM180327 | Procell |
| Trypsin-EDTA | C0201 | Beyotime |
| Lipo8000™ transfection reagent | C0533 | Beyotime |
| HiScript III RT SuperMix (qPCR) | R333 | Vazyme |
| Advanced DMEM/F12 | 12634-010 | Gibco |
| A83-01 | SML0788 | Sigma |
| B-27supplement, serum free | 17504044 | Gibco |
| Collagenase, type II | 17101015 | Gibco |
| DNase I | 11284932001 | Roche |
| GlutaMAX supplement | 35050061 | Gibco |
| HEPES | 15630080 | Gibco |
| Human Noggin | 120-10C | Peprotech |
| Liberase™ | 5401119001 | Sigma |
| Matrigel growth factor reduced (GFR) basement membrane matrix | 356231 | Corning |
| N-Acetylcysteine | A9165 | Sigma |
| Nicotinamide | N0636 | Sigma |
| Recombinant human FGF-10 | BY0017 | Biodragon |
| Recombinant human KGF (FGF-7) | BY0015 | Biodragon |
| Recombinant human R-spondin 1 protein | BY0049 | Biodragon |

| Culture material/Reagent/Kit Name | Catalog Number | Source |
|---|---|---|
| SB202190 Monohydrochloride hydrate | S7076 | Sigma |
| TrypLE Express | 12605-010 | Gibco |
| Y-27632 dihydrochloride | M1817 | Abmole Bioscience |
| Taq Pro SYBR qPCR Master Mix | Q712 | Vazyme |
| Antifade Mounting Medium with DAPI | P0131 | Beyotime |
| Phosphatase inhibitor cocktail | P1260 | Solarbio |
| ECL substrate | P0018FS | Beyotime |
| Protein A+G Agarose beads | P2055 | Beyotime |
| BCA protein assay kit | P0009 | Beyotime |
| Enhanced BCA Protein Assay Kit | P0009 | Beyotime |
| Dual-Luciferase Reporter Assay Kit | RG027 | Beyotime |
| Plasmid extraction kit | D6943-01 | Omega |
| TUNEL Assay Kit | BR3108871 | Bioleaper |
| BeyoChIP™ Enzymatic ChIP Assay Kit | P2083S | Beyotime |
| Organoid Vitality Assay Kit | HY-K6016 | MCE |
